# Fine-tuning GPCR-mediated neuromodulation by biasing signaling through different G-protein subunits

**DOI:** 10.1101/2023.03.03.529094

**Authors:** Jong-Chan Park, Alex Luebbers, Maria Dao, Ana Semeano, Maria P. Papakonstantinou, Stefan Broselid, Hideaki Yano, Kirill A. Martemyanov, Mikel Garcia-Marcos

**Affiliations:** Department of Biochemistry, Chobanian and Avedisian School of Medicine, Boston University, Boston, MA 02118, USA; UF Scripps Biomedical Research, University of Florida, Jupiter, FL 33458; Department of Pharmaceutical Sciences, Center for Drug Discovery, School of Pharmacy and Pharmaceutical Sciences, Bouvé College of Health Sciences, Northeastern University, Boston MA 02115; Department of Biology, College of Arts and Sciences, Boston University, Boston, MA 02115, USA

**Author notes:** These authors contributed equally to this work (Jong-Chan Park, Alex Luebbers). Corresponding author: Mikel Garcia-Marcos.

## Abstract

GPCRs mediate neuromodulation through activation of heterotrimeric G-proteins (Gαβγ). Classical models depict that G-protein activation leads to a one-to-one formation of Gα-GTP and Gβγ species. Each of these species propagates signaling by independently acting on effectors, but the mechanisms by which response fidelity is ensured by coordinating Gα and Gβγ responses remain unknown. Here, we reveal a paradigm of G-protein regulation whereby the neuronal protein GINIP biases inhibitory GPCR responses to favor Gβγ over Gα signaling. Tight binding of GINIP to Gαi-GTP precludes its association with effectors (adenylyl cyclase) and, simultaneously, with Regulator-of-G-protein-Signaling (RGS) proteins that accelerate deactivation. As a consequence, Gαi-GTP signaling is dampened whereas Gβγ signaling is enhanced. We show that this mechanism is essential to prevent imbalances of neurotransmission that underlie increased seizure susceptibility *in vivo*. Our findings reveal an additional layer of regulation within a quintessential mechanism of signal transduction that sets the tone of neurotransmission.

## INTRODUCTION

Understanding the molecular basis for how neurotransmission is orchestrated remains a central challenge in neuroscience. Dissecting the mechanisms by which hundreds of neurotransmitter receptors propagate extracellular chemical signals to the interior of neurons is of utmost importance for the control of neuronal excitability. While ionotropic receptors, which are ligand-gated ion channels, directly regulate ion fluxes, metabotropic receptors affect ionic activity indirectly through second messengers and other signaling intermediaries. G protein-coupled receptors (GPCRs) are the largest family of metabotropic receptors, which collectively mediate responses to most neurotransmitters (Gilman, 1987; Roth, 2019; Weis and Kobilka, 2018). They can be found at pre-synaptic and post-synaptic structures, where they modulate synaptic transmission in a “slow” time scale of seconds to minutes due to the requirement of signaling intermediaries like heterotrimeric G-proteins (Gαβγ) (Greengard, 2001; Roth, 2019; Zurawski et al., 2019). GPCRs are also the largest class of druggable targets in the human genome. They represent the target for >30% of drugs approved for clinical use, including numerous medications for neurological and neuropsychiatric diseases (Hauser et al., 2017; Hopkins and Groom, 2002; Santos et al., 2017; Sriram and Insel, 2018). However, the molecular mechanisms that control GPCR-mediated neuromodulation are very complex and remain the subject of intense investigation.

The main mechanism of action of GPCRs is through activation of heterotrimeric G-proteins, although some responses are also mediated via β-arrestins (Weis and Kobilka, 2018). GPCRs activate heterotrimeric G-proteins (Gαβγ) by promoting GTP loading on Gα. Then, Gα-GTP and Gβγ dissociate (or rearrange), which allows their binding to and modulation of downstream effectors to propagate signaling. Signaling is turned off upon GTP hydrolysis, leading to re-association of Gα with Gβγ. While this mechanism of G-protein regulation by GPCRs is well understood, much less is known about how it is influenced by a growing body of cytoplasmic proteins that act on G-proteins (Cismowski et al., 1999; De Vries et al., 1995; De Vries et al., 2000b; DiGiacomo et al., 2018; Dohlman and Thorner, 1997; Ross and Wilkie, 2000; Sato et al., 2006; Siderovski and Willard, 2005). Most, if not all, of these regulators have in common that they bind to Gα subunits and affect their enzymatic activity by modulating nucleotide binding or hydrolysis, thereby having a profound impact on the duration and amplitude of G-protein signaling. For example, Regulators of G-protein Signaling (RGS) are <u>G</u>TPase <u>A</u>ctivating <u>P</u>roteins (GAPs) that bind to Gα-GTP and accelerate the rate of nucleotide hydrolysis, which facilitates G-protein signaling termination (Berman et al., 1996b; Watson et al., 1996). Other cytoplasmic regulators alter nucleotide exchange rates (i.e., GTP loading) by serving as <u>G</u>uanine-nucleotide <u>D</u>issociation Inhibitors (GDIs) (De Vries et al., 2000a; Kimple et al., 2001; Peterson et al., 2000) or as non-receptor <u>G</u>uanine-nucleotide <u>E</u>xchange <u>F</u>actors (GEFs) (Cismowski et al., 2000; DiGiacomo et al., 2018; Lee and Dohlman, 2008; Tall et al., 2003).

Heterotrimeric G-proteins are broadly divided in four families based on sequence and functional similarity (G_s_, G_i/o_, G_q/11_, G_12/13_) (Gilman, 1987). In the nervous system, G_i/o_ proteins mediate GPCR-mediated inhibitory neuromodulation triggered upon stimulation by neurotransmitters like GABA, dopamine, serotonin, or norepinephrine, among many others. When activated, G_i/o_ proteins release free Gβγ subunits that modulate numerous targets, including inhibition of pre-synaptic voltage-gated Ca^2+^ channels (e.g., Ca_v_2) or activation of post-synaptic G protein-gated inwardly rectifying K^+^ channels (GIRK also known as Kir3) (Betke et al., 2012; Csanady, 2017). These direct effects on ion activity suppress neurotransmission by preventing neurotransmitter release or reducing post-synaptic excitation, respectively. At the same time, G_i/o_ activation can also lead to the formation of Gαi-GTP, which inhibits adenylyl cyclases (Ostrom et al., 2022; Sadana and Dessauer, 2009). This dampens the intracellular levels of cAMP, which has indirect effects on neurotransmission through the numerous targets affected by this ubiquitous second messenger (Hosaka et al., 1999; Kandel, 2012; Lonart et al., 2003; Nagy et al., 2004; Westphal et al., 1999). Given the large complexity of signaling networks affected simultaneously by Gβγ and Gα-GTP, a central and unresolved question in GPCR signaling is: how is response specificity achieved? In particular, the fact that Gβγ and Gα-GTP are generated at equimolar amounts upon GPCR stimulation poses the conundrum of how different signaling cascades triggered by each one of the two species independently are coordinated to ensure the fidelity of the cellular response.

Here, we characterize a mechanism for biasing G_i_ responses in which Gαi-GTP signaling is inhibited while, *simultaneously*, Gβγ signaling is enhanced after receptor stimulation. We demonstrate that this mechanism of signaling coordination is required to ensure appropriate scaling of GPCR neuromodulatory influence over neuronal excitability. More specifically, our findings indicate that GINIP (aka KIAA1045), a protein previously identified as a binder of active Gαi (Gaillard et al., 2014; Maziarz et al., 2018) and as a regulator of GABA_B_ receptor (GABA_B_R) signaling in dorsal root ganglia (Gaillard et al., 2014; Liu et al., 2016), is a broad regulator of signaling via G_i_-coupled GPCRs expressed across multiple brain regions in both inhibitory and excitatory neurons. At the molecular level, GINIP biases receptor responses to favor Gβγ signaling over Gαi-GTP signaling without affecting the enzymatic activity of Gα subunits directly, which contrasts with the mechanisms of action of other G-protein regulators described to date. Overall, our findings reveal an additional layer of GPCR regulation through a previously unknown form of biased signaling that has a crucial role in modulating neurotransmission.

## RESULTS

### GINIP binds to active Gαi subunits with high affinity

A prior report showed that GINIP binds to GTPase-deficient Gαi mutants that mimic active G proteins (Gaillard et al., 2014), but no rigorous biochemical characterization was performed. Using purified GINIP and Gαi3 proteins, we found that GINIP binds with high affinity (K_D_ ~70 nM) to *bona fide* active Gαi3 (i.e., loaded with GTPγS, a non-hydrolysable GTP analog), as well as to the active transition state for GTP hydrolysis adopted upon binding of GDP-AlF_4^−^_ (Coleman et al., 1994; Tesmer et al., 1997a) (**Fig. 1A**). Binding to inactive, GDP-bound Gαi3 was undetectable (**Fig. 1A**). Further protein binding experiments revealed that GINIP binds similarly to all three Gαi isoforms (Gαi1, Gαi2 and Gαi3), but not to other α-subunits of the G_i/o_ family like Gαo and Gαz, or to representative members of any of the other three G-protein families (i.e., Gαs for G_s_, Gαq for G_q/11_, or Gα12 for G_12/13_) (**Fig. 1B**, **Fig. S1A**). These results indicate that GINIP is highly selective for Gαi isoforms, and that it binds with a marked G-protein state preference for active Gα.

**Figure 1.**
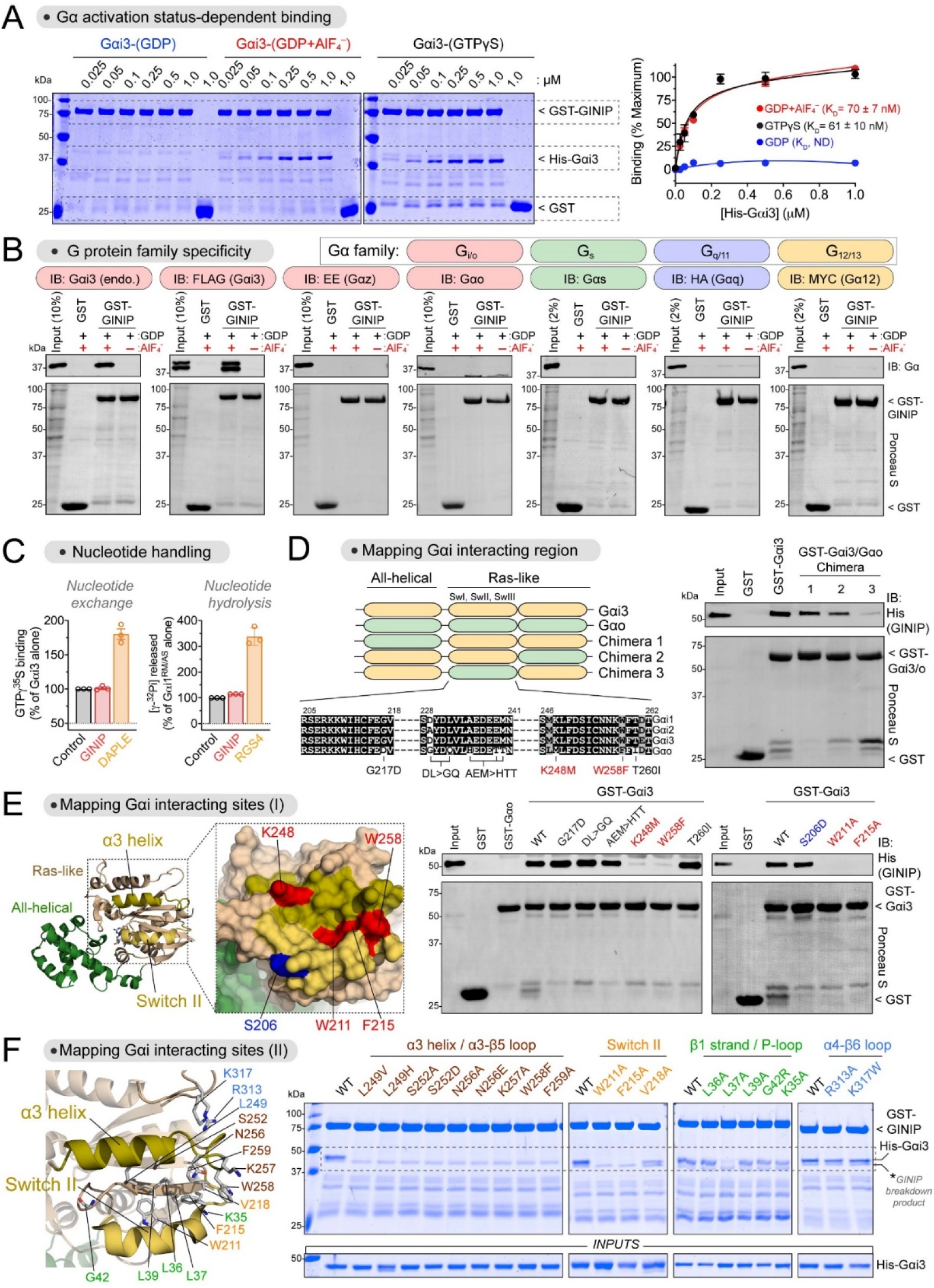
GINIP binds to the effector binding region of Gαi without affecting its enzymatic activity. **(A)** GINIP binds to active but not inactive Gαi3. *Left*, representative Coomassie-stained gel showing binding of His-Gαi3 loaded with the indicated nucleotides (GDP, GDPꞏAlF_4^−^_, GTPγS) to immobilized GST-GINIP. *Right*, quantification of His-Gαi3 binding to GST-GINIP based on band densitometry. K_D_’s were determined based on fitting the data to a one-site binding nonlinear regression curve. Mean ± S.E.M., n=3-4. **(B)** GINIP binds to Gαi3, but not to other Gα’s of the same family (Gαo, Gαz), or other families (Gαs, Gαq, Gα12). Lysates of HEK293T cells expressing the indicated G proteins were incubated with GST or GST-Gαi3 immobilized on glutathione-agarose beads in the presence of GDP or GDPꞏAlF_4^−^_, as indicated. Bead-bound proteins were detected by Ponceau S staining or by immunoblotting (IB). **(C)** GINIP does not affect the enzymatic activity of Gαi. Nucleotide exchange on Gαi3 was determined by GTPγS binding, whereas nucleotide hydrolysis by Gαi1^RM/AS^ was determined by the production of free phosphate (Pi) from GTP. GINIP, 2 μM, DAPLE, 1 μM, RGS4, 0.2 μM. Mean ± S.E.M., n=3. **(D)** Gαi3 region 205-262 is required for GINIP binding. *Left*, diagram of Gαi3 (orange) /Gαo (green) chimeras. Sequence alignment of the Gαi3 205-262 region with Gαi1, Gαi2 and Gαo, indicating mutations tested in panel (E). *Right*, representative experiment showing binding of purified His-GINIP to the indicated G proteins in the presence of GDPꞏAlF_4^−^_. Bead-bound proteins were detected by Ponceau S staining or by immunoblotting (IB). **(E)** Mutation of residues in the α3 helix and Switch II of Gαi ablate GINIP binding. *Left,* Structural model of Gαi1-(GDPꞏAlF_4^−^_) (PDB: 2G83). Red indicates residues in the α3/Switch II region that result in loss of GINIP when mutated, whereas blue indicates a residue in the same region that does not affect GINIP binding when mutated. *Center,* Mutation of some residues in the α3 helix from Gαi3 to cognate amino acids from Gαo ablate binding to GINIP. *Right,* representative experiment showing binding of purified His-GINIP to the indicated G proteins in the presence of GDPꞏAlF_4^−^_. Bead-bound proteins were detected by Ponceau S staining or by immunoblotting (IB). **(F)** Mutation of residues within, but not adjacent to, the effector binding region (α3/Switch II groove) of Gαi impair GINIP binding. *Left*, Structural model of Gαi1-(GDPꞏAlF_4^−^_) (PDB: 2G83) displaying the residues investigated by site-directed mutagenesis. *Right*, representative Coomassie-stained gel showing binding of the indicated Gαi3 proteins loaded with GTPγS to immobilized GST-GINIP. All protein electrophoresis results are representative of n ≥ 3 experiments.

### GINIP does not affect the enzymatic activity of Gαi subunits

Proteins that bind Gα in a state-dependent manner tend to either serve as effectors or regulate G-proteins by modulating their ability to bind or hydrolyze nucleotides (Cismowski et al., 2000; DiGiacomo et al., 2018; Dohlman and Thorner, 1997; Lee and Dohlman, 2008; Ross and Wilkie, 2000; Sato et al., 2006; Siderovski and Willard, 2005; Sunahara et al., 1997; Tall, 2013). Since GINIP lacks any known catalytic domain that could justify its ability to propagate signaling by serving as a G-protein effector, we investigated if GINIP affects nucleotide handling by Gαi subunits. First, we found that nucleotide exchange by Gαi3, as determined by an established GTPγS binding assay (Garcia-Marcos et al., 2010; Leyme et al., 2014a), was not affected by GINIP, whereas the guanine-nucleotide exchange factor DAPLE (Aznar et al., 2015) led to an increase in activity under the same conditions (**Fig. 1C**). Next, we determined if GINIP affects nucleotide hydrolysis by Gαi. For this, we did steady-state GTPase assays using a Gαi1 mutant (Gαi1^RM/AS^) for which nucleotide hydrolysis is the rate limiting step under these conditions (Zielinski et al., 2009). We found that GINIP did not affect nucleotide hydrolysis by Gαi1^RM/AS^, despite binding to it as strongly as to Gαi1 wild-type (**Fig. S1A**), whereas the GAP RGS4 (Berman et al., 1996b) led to a marked increase in activity (**Fig. 1C**). Taken together, these results show that GINIP does not affect the enzymatic activity of Gαi despite binding to it with high affinity.

### GINIP binds to the groove formed by the α3 helix and the switch II region of Gαi

To begin elucidating the functional consequences of GINIP binding to Gαi, we carried out experiments to map its binding site on the G-protein with the hope that structure would inform about function. First, we leveraged the marked preference of GINIP for binding Gαi isoforms over the closely related protein Gαo (**Fig. 1B**, **Fig.S1A**) to design G-protein chimeras (**Fig. 1D**). We generated three Gαi/o chimeras by replacing, one at a time, three different regions of Gαi3 with the corresponding regions of Gαo and performed GINIP binding experiments with them. We found that the region between amino acid (aa) 178 and aa 270 of Gαi3 was required for GINIP binding (**Fig. 1D**). This segment is located in the Ras-like domain of Gα and encompasses the three “switch” regions that change conformation between GDP- and GTP-bound states (Coleman et al., 1994; Mixon et al., 1995), as well as other elements like the α3 helix. Next, we identified specific positions in the aa178-270 region bearing residues that are different between Gαo and Gαi subunits, and systematically replaced them in Gαi3 for their counterparts in Gαo (**Fig. 1D, E**). Of these, K248M and W258F completely abolished GINIP binding to active Gαi3 (**Fig. 1E**). K248 is located in the α3 helix and W258 in the adjacent α3/β5 loop. In the crystal structure of active Gαi (Coleman et al., 1994; Tesmer et al., 1997a), both residues are oriented towards a groove formed between the α3 helix and the switch II (SwII) region (**Fig. 1E**), which is a pocket utilized by Gα subunits to bind effectors and effector-like interactors (Chen et al., 2005; Johnston et al., 2006; Slep et al., 2001; Tesmer et al., 1997b; Tesmer et al., 2005; Waldo et al., 2010). Mutation of two SwII residues (W211 and F215) that contribute to this pocket also disrupted GINIP binding, whereas mutation of another SwII residue not oriented towards the α3/SwII groove (S206) did not (**Fig. 1E**).

We further pinpointed the structural determinants of Gαi3 required for GINIP binding by using a battery of mutants spanning regions within or adjacent to the α3/SwII groove (**Fig. 1F**). Of these, mutation of residues in the α3 helix (L249, S252, N256), α3/β5 loop (K257, W258, F259) and SwII (W211, F215, V218) oriented towards the α3/SwII groove resulted in loss of GINIP binding to active Gαi3 (**Fig. 1F**). Moreover, mutation of a residue of the β1 strand positioned at the bottom of the α3/SwII groove (L37) also resulted in loss of GINIP binding (**Fig. 1F**). In contrast, mutation of other residues adjacent to the elements forming the α3/SwII groove did not disrupt GINIP binding (K35, L36, L38, G42 in the β1 strand/P-loop, or R313, K317 in the α4/β6 loop, **Fig. 1F**). All mutants used in these experiments have been previously shown to be functional based on their ability to adopt an active conformation upon GTP binding (de Opakua et al., 2017; Garcia-Marcos et al., 2010). These results indicate that residues in the α3/SwII groove of Gαi are specifically involved in mediating GINIP binding. Consistent with this idea, we also found that GINIP competes for binding to Gαi3 with the effector-like peptide KB-1753, which binds on the α3/SwII groove (Johnston et al., 2006) (**Fig. S1B**). Taken together, these results provide strong evidence that the α3/SwII groove of Gαi is a critical binding site for GINIP.

### GINIP dampens cAMP inhibition triggered by multiple G_i_-coupled GPCRs

G_i_ proteins (Gαiꞏβγ) were discovered based on and are defined by their ability to inhibit the activity of adenylyl cyclases, which are the canonical effectors for Gαi-GTP (Ostrom et al., 2022; Sadana and Dessauer, 2009). Based on our finding that GINIP binds with high affinity to the effector binding site of Gαi-GTP, we hypothesized that GINIP would prevent the action of the G-protein on adenylyl cyclase. If so, GINIP should dampen cAMP inhibition upon G_i_ activation regardless of the GPCR responsible for triggering such activation. Consistent with a previous report (Gaillard et al., 2014), we found that GINIP reduced the extent by which the GABA_B_R inhibited forskolin-stimulated cAMP in HEK293T cells (**Fig. 2A**). Moreover, similar results were obtained with two other unrelated neurotransmitter GPCRs that activate G_i_, the α2_A_ adrenergic receptor (α2_A_-AR) and the D2 dopamine receptor (D2R) (**Fig. 2A**). In contrast, GINIP expression did not affect forskolin-stimulated cAMP accumulation in the same assay format (0.66 ± 0.10 vs 0.60 ± 0.07 ΔBRET^−1^ in the presence or absence of GINIP expression, respectively, 10 min after stimulation with 1 μM forskolin, n=8, p>0.05, paired t-test), indicating that GINIP has no direct effect on adenylyl cyclase activity. Overall, these results support the idea that the effects of GINIP occur at the level of the G-protein rather than being receptor specific.

**Figure 2.**
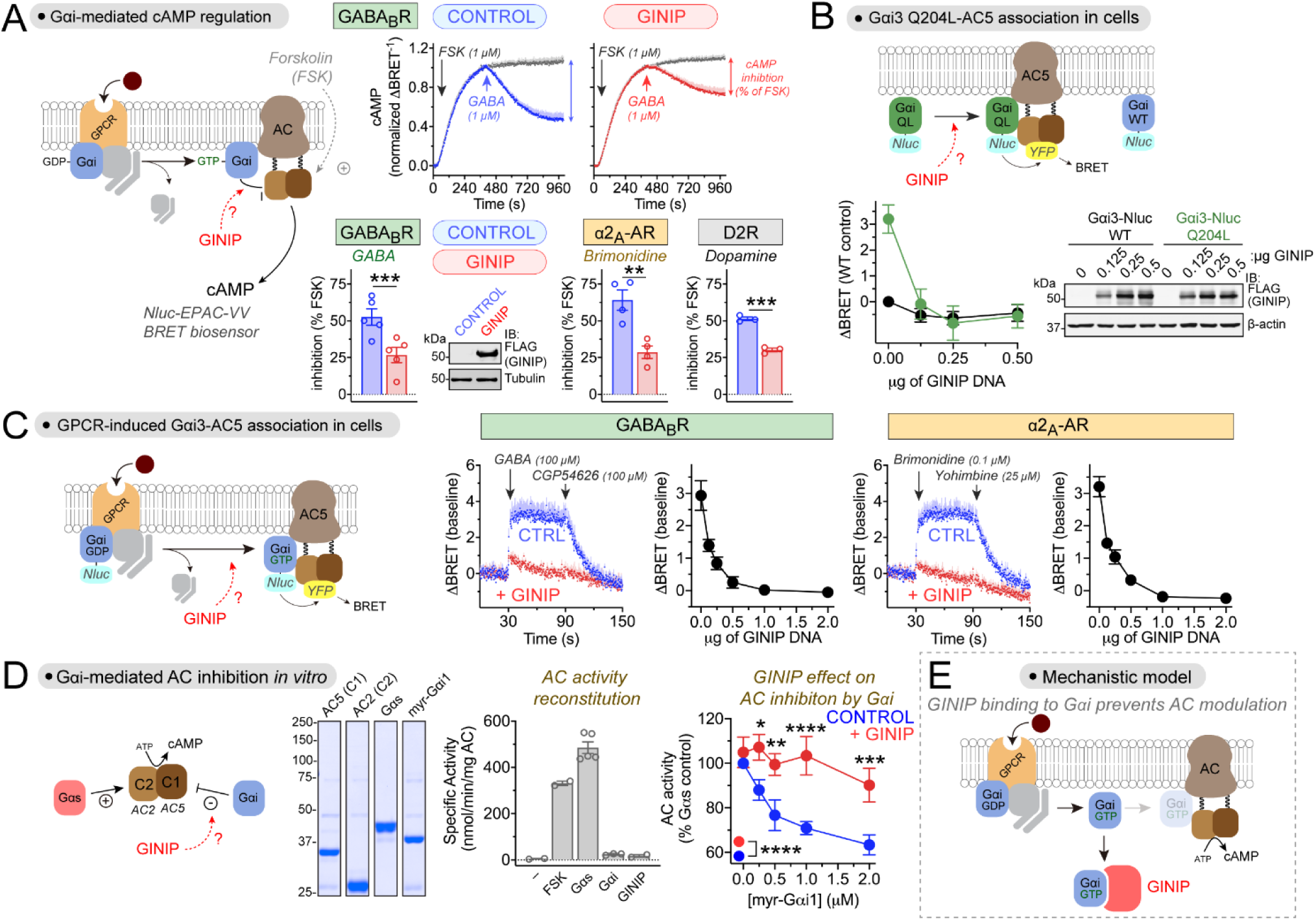
GINIP directly blocks Gαi-mediated regulation of adenylyl cyclase. **(A)** GINIP prevents Gαi-mediated inhibition of adenylyl cyclase (AC) upon stimulation of 3 different G_i_-coupled GPCRs. *Top row,* kinetic traces of BRET measurement of cAMP in HEK293T cells expressing the GABA_B_R in the presence or absence of GINIP treated with forskolin (FSK) and GABA as indicated. *Bottom row,* quantified inhibition of FSK-stimulated cAMP upon stimulation of GABA_B_R, α2-AR, or D2R with GABA (1 μM), Brimonidine (5 μM), or dopamine (0.2 μM). Immunoblot (IB) validates GINIP expression. Mean ± S.E.M., n=3-5. **p<0.01, ***p<0.001, paired t-test. **(B)** GINIP prevents the association of active Gαi3 with AC5 in cells. *Left,* changes in BRET (ΔBRET) were determined in HEK293T cells expressing Gαi3-Nluc WT or Gαi3-Nluc Q204L upon transfection of increasing amounts of GINIP. Mean ± S.E.M., n=6. *Right*, validation of GINIP expression by IB. **(C)** GINIP prevents the association of Gαi3 with AC5 upon GPCR stimulation. BRET was measured in HEK293T cells expressing the GABA_B_R or the α2_A_-AR upon transfection of different amounts of GINIP DNA. Kinetic traces correspond to cells expressing no GINIP (‘CTRL’ blue) or transfected with 2 μg of GINIP plasmid (red). Cells were treated with the indicated GPCR agonists/antagonists, and BRET changes (ΔBRET) one minute after agonist stimulation for each amount of GINIP transfection. Mean ± S.E.M., n=3. **(D)** GINIP blocks the regulation of AC by Gαi *in vitro*. Coomassie-stained gel shows the purified proteins used to reconstitute and modulate AC activity *in vitro*. Bar graph shows that FSK (5 μM) or Gαs-GTPγS (0.1 μM), but not myr-Gαi1 (2 μM) or GINIP (2 μM), promote the activation of reconstituted AC (AC5 (c1) + AC2 (C2)). *Right,* Gαs-stimulated AC activity was determined in the presence of increasing concentrations of myr-Gαi1-GTPγS in the absence (blue) or presence of GINIP (2 μM, red). Mean ± S.E.M., n=5. *p<0.05, **p<0.01, ***p<0.001, ****p<0.0001, two-way ANOVA for presence/ absence of GINIP x myr-Gαi1 concentration, with multiple comparisons at each concentration using Fisher’s LSD test. **(E)** Diagram summarizing the proposed mechanism action of GINIP on Gαi-mediated modulation of AC activity (competitive binding).

### GINIP prevents the association of Gαi with adenylyl cyclase in cells

To further test the hypothesis that GINIP regulates cAMP responses by preventing the association between Gαi-GTP and effectors, we used a bioluminescence resonance energy transfer (BRET) assay to monitor the interaction between Gαi3 and adenylyl cyclase 5 (AC5) in HEK293T cells (**Fig. 2B, C**). For this, Gαi3 internally tagged with nanoluciferase (Nluc) was co-expressed in cells with YFP-tagged AC5. First, we observed that a constitutively active mutant of Gαi3 (Gαi3 Q204L) that mimics Gα-GTP resulted in higher BRET levels than Gαi3 wild-type (WT) in the absence of GPCR stimulation (**Fig. 2B**), which is consistent with the expected preferential binding of AC5 with active G-proteins. Co-expression of increasing amounts of GINIP resulted in a decrease of BRET in cells expressing Gαi3 Q204L to levels equivalent to those observed for Gαi3 WT (**Fig. 2B**), indicating that GINIP prevents the association of active G-proteins with adenylyl cyclase. Next, we used the same BRET assay to investigate the effect of GINIP on GPCR-stimulated Gαi3-AC5 association (**Fig. 2C**). We found that expression of increasing amounts of GINIP efficiently suppressed BRET responses induced by the stimulation of either GABA_B_R or α2_A_-AR (**Fig. 2C**). These results not only indicate that GINIP prevents the association of Gαi with its effector adenylyl cyclase, but also that this effect is independent of the GPCR triggering G_i_ activation.

### GINIP inhibits directly the regulation adenylyl cyclase by Gαi-GTP

To more definitely pinpoint that the mechanism of action of GINIP is direct blockade of Gαi-effector interaction, we used a reductionist approach with G-proteins and adenylyl cyclase as purified components in the presence or absence of purified GINIP. We reconstituted adenylyl cyclase activity *in vitro* with the purified C1 domain of AC5 and the purified C2 domain of AC2, as previously described (Sunahara et al., 1997), which led to robust cAMP synthesis when incubated together in the presence of forskolin or purified Gαs loaded with GTPγS (**Fig. 2D**). Consistent with previous observations (Dessauer et al., 1998), purified myristoylated Gαi1 (myr-Gαi1) loaded with GTPγS inhibited Gαs-stimulated adenylyl cyclase activity in a dose-dependent manner. This inhibitory effect of myr-Gαi1 was suppressed in the presence of GINIP (**Fig. 2D**), demonstrating that GINIP directly prevents the regulation of adenylyl cyclase by Gαi.

Taken together, our results support a mechanism in which GINIP regulates GPCR-dependent signaling by directly binding to Gαi-GTP and preventing its association with the effector adenylyl cyclase, therefore resulting in the suppression of Gα-GTP dependent signaling (**Fig. 2E**).

### GINIP promotes free Gβγ signaling upon GPCR stimulation

After establishing how GINIP affects Gαi-GTP dependent signaling, we turned our attention to Gβγ, the other signaling species that emerges from GPCR-mediated activation of heterotrimeric G-proteins. For this, we leveraged a BRET-based biosensor that directly detects levels of “free” Gβγ (i.e., Gβγ competent for engaging its effectors) (Hollins et al., 2009; Masuho et al., 2015) (**Fig. 3A**). We found that expression of GINIP in HEK293T cells increased the amplitude of Gβγ responses elicited upon stimulation of GABA_B_R with its agonist GABA at a submaximal concentration (1 μM) but not at a maximal concentration (100 μM) (**Fig. 3A**). However, when we determined the rate of Gβγ deactivation upon addition of an antagonist post-GABA stimulation, we found that GINIP delayed the rate of deactivation regardless of the concentration of agonist used (**Fig. 3A**). The magnitude of the change in deactivation rate was dependent on the amount of GINIP expressed (**Fig. S2A, B**). Similar effects of GINIP were observed with a second G_i_-coupled GPCR, the α2_A_-AR (**Fig. S2C**), suggesting that GINIP acts at the level of the G-protein instead of being receptor-specific. These results indicate that the overall effect of GINIP on Gβγ signaling is to enhance, rather than suppress as observed for Gαi-GTP signaling.

**Figure 3.**
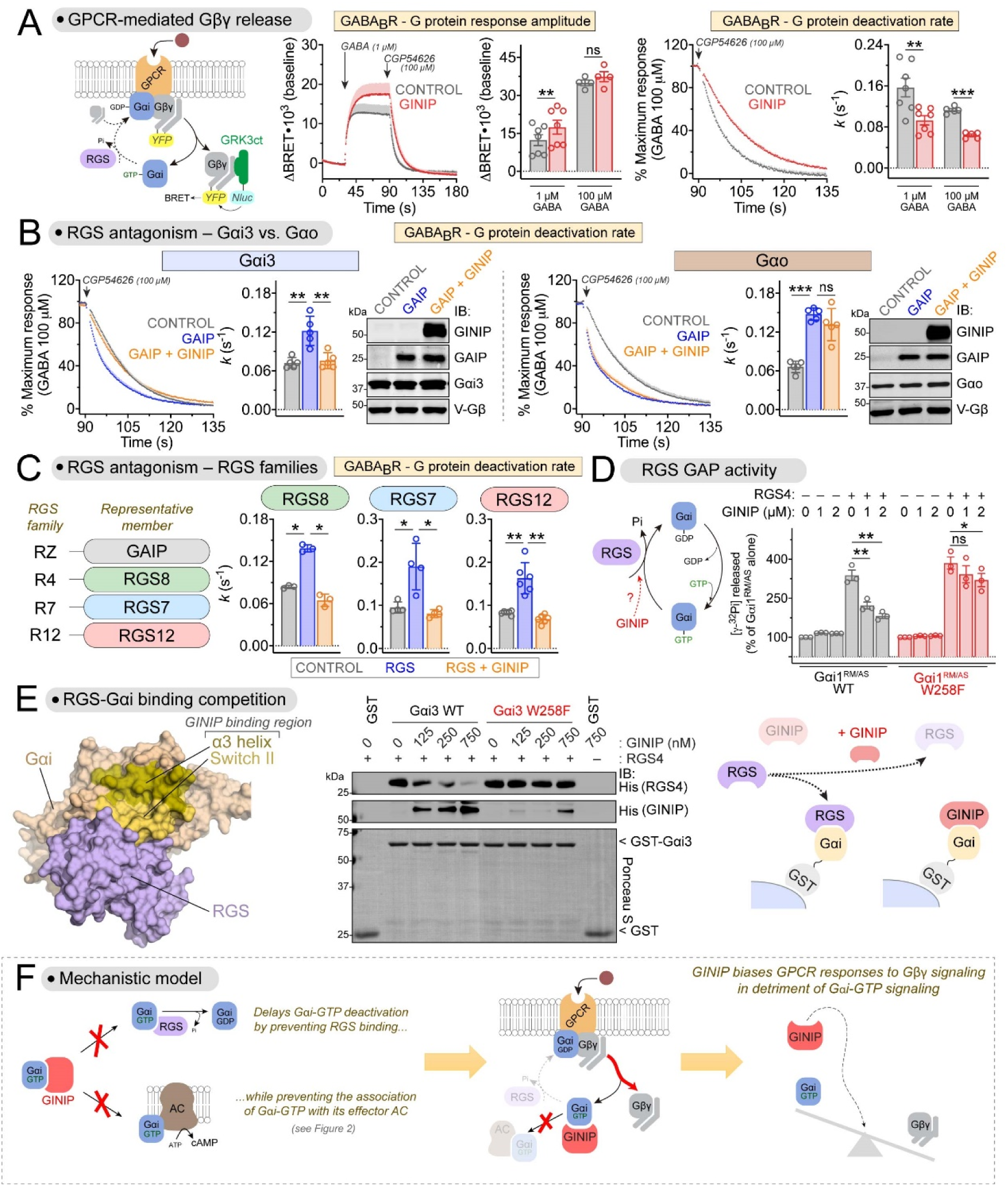
GINIP promotes Gβγ-mediated signaling by antagonizing the action of RGS GAPs on Gαi. **(A)** GINIP enhances Gβγ-mediated signaling triggered by GABA_B_R. *Left,* diagram of G protein activation/deactivation cycle and BRET-based detection of free Gβγ. *Center,* BRET was measured in HEK293T cells expressing the GABA_B_R in the absence (black) or presence (red) of GINIP. Kinetic traces correspond to cells expressing no GINIP (black) or transfected with 2 μg of GINIP plasmid (red). Cells were treated with GABA and CGP54626 as indicated and the amplitude of the BRET responses quantified 1 min after agonist stimulation. *Right,* G protein deactivation rates were determined by normalizing the BRET data to maximum response and fitting the post-antagonist data to an exponential decay curve to extract rate constant values (*k*). Mean ± S.E.M., n=4-7. ns = not significant, **p<0.01, ***p<0.001, paired t-test. **(B)** GINIP antagonizes GAIP-mediated acceleration of Gβγ deactivation for Gi but not Go proteins. BRET experiments were carried out and analyzed as in (A) with cells expressing Gαi3 or Gαo in the absence (grey) or presence of GAIP (blue) or GAIP plus GINIP (orange). Expression of GAIP and GINIP was validated by immunoblotting IB. Mean ± S.E.M., n=5. ns = not significant, **p<0.01, ***p<0.001, one-way ANOVA corrected for multiple comparisons (Tukey). **(C)** GINIP antagonizes the acceleration of Gβγ deactivation mediated by representative members of all RGS families. BRET experiments were carried out and analyzed as in (B), expect that RGS8 (R4), RGS7 (R7), or RGS12 (R12) were used instead of GAIP (RZ). For experiments with RGS7, cells were also co-transfected with plasmids encoding Gβ_5_ and R7BP. Mean ± S.E.M., n=3-6. *p<0.05, **p<0.01, one-way ANOVA corrected for multiple comparisons (Tukey). **(D)** GINIP antagonizes the GAP activity of RGS4 on Gαi *in vitro*. Nucleotide hydrolysis by Gαi1^RM/AS^ (WT or W258F) was determined in the presence of RGS4 and/or GINIP as indicated. Mean ± S.E.M., n=3. ns = not significant, *p<0.05, **p<0.01, one-way ANOVA corrected for multiple comparisons (Tukey). **(E)** GINIP competes with RGS4 for binding to Gαi3. *Left,* Structural model of Gαi1-(GDPꞏAlF_4^−^_) bound to RGS4 (PDB: 1AGR). *Right,* increasing concentrations of purified His-GINIP and a fixed amount of His-RGS4 (20 nM) were incubated with GST or GST-Gαi3 (WT or W258F) immobilized on glutathione-agarose beads in the presence of GDPꞏAlF_4^−^_. Bead-bound proteins were detected by Ponceau S staining or by immunoblotting (IB). One representative result of three independent experiments is shown. **(F)** Diagram summarizing the proposed mechanism by which GINIP biases G protein responses by favoring Gβγ-dependent signaling in detriment of Gαi-dependent signaling. Binding of GINIP to G proteins simultaneously impairs Gαi binding to its effector (AC) and to negative regulators (RGS GAPs).

### GINIP antagonizes RGS GAP-mediated deactivation of Gβγ signaling

The effects of GINIP on Gβγ signaling are reminiscent of those observed when the function of RGS proteins is disabled (Lambert et al., 2010). RGS proteins are GAPs for Gα subunits that negatively regulate GPCR responses (Dohlman et al., 1996; Druey et al., 1996; Kimple et al., 2011). When RGS protein function is disabled, Gβγ deactivation rates slow down and response amplitudes are potentiated (Lambert et al., 2010), much like what we found upon GINIP expression (**Fig. 3A**). Based on this, we hypothesized that GINIP might antagonize the action of RGS GAPs. Expression of the RGS protein GAIP (aka RGS19) accelerated the rate of Gβγ deactivation upon modulation of G_i3_ protein activity by GABA_B_R, as expected, and this effect was suppressed when GINIP was co-expressed with GAIP (**Fig. 3B**). Similar results were obtained with the α2_A_-AR (**Fig. S2D**), suggesting that GINIP antagonizes the inhibitory action of RGS proteins on Gi proteins independent of the receptor responsible for their activation. In contrast, GINIP did not antagonize the acceleration of Gβγ deactivation by GAIP when the G-protein modulated by GABA_B_R was G_o_ instead of G_i3_ (**Fig. 3B**). These results are consistent with the lack of binding of GINIP to Gαo (**Fig. 1**), and further confirm that GINIP exerts its effects on Gβγ signaling by acting at the level of the Gα subunit rather than any other component of the signaling system. We also assessed if the effect of GINIP on GAIP-mediated regulation of Gβγ signaling was a phenomenon generalizable to other RGS proteins, with the expectation that it would be because all RGS proteins share a similar binding mode and mechanism of action on Gα (Soundararajan et al., 2008; Tesmer et al., 1997a). In addition to the RZ family to which GAIP belongs, there are 3 other RGS proteins families: R4, R7 and R12. Our results confirmed that GINIP can suppress the acceleration of Gβγ deactivation exerted by representative members of each of these families— i.e., RGS8 (R4), RGS7 (R7), or RGS12 (R12) (**Fig. 3C**). These findings show that GINIP antagonizes the regulation of Gβγ signaling by all four RGS protein families.

### GINIP inhibits directly the regulation of Gαi by RGS proteins

Since GINIP binds directly to Gαi (**Fig. 1**) and the cell-based experiments above indicate RGS protein antagonism at the level of Gα (**Fig. 3B, C**), we tested if GINIP directly inhibits the GAP activity of RGS proteins by using purified proteins. We found that GINIP inhibited the GAP activity of RGS4 or GAIP, as determined by their ability to enhance the rate of nucleotide hydrolysis by Gαi1^RM/AS^ (**Fig. 3D**, **Fig. S3A, B, D**). In contrast, this effect of GINIP was not observed when the G-protein contained the W258F mutation (**Fig. 3D**, **Fig. S3D**) that precludes GINIP binding (**Fig. 1**) while preserving Gαi binding to RGS proteins (**Fig. S3C** and (Garcia-Marcos et al., 2010)). These results demonstrate that GINIP blocks the GAP activity of RGS proteins by binding to Gαi.

### GINIP competes with RGS proteins for binding to Gαi

We reasoned that the mechanism by which GINIP blocks RGS GAP activity is competition for Gαi binding. This is not only because disrupting GINIP binding to Gαi prevents the inhibition of RGS GAP activity (**Fig. 3D**, **Fig. S3D**), but also because of the following two features of how GINIP binds Gαi: (1) GINIP has high affinity for the same conformation that is preferred by RGS proteins (**Fig. 1A**), i.e., the transition state mimicked by GDP-AlF_4^−^_ activation (Berman et al., 1996a; Tesmer et al., 1997a), and (2) it binds at a site (α3/SwII groove, **Fig. 1**) that is adjacent to the RGS binding region (**Fig. 3E**, *left*). To test this mechanism, we determined RGS4 binding to Gαi3 loaded with GDP-AlF_4^−^_ in the absence or presence of GINIP using purified proteins (**Fig. 3E**). We found that binding of GINIP to Gαi3-GDP-AlF_4^−^_ was accompanied by a marked reduction of RGS4 binding, which became barely detectable at the highest concentration of GINIP tested (**Fig. 3E**). In contrast, GINIP did not affect binding of RGS4 to the Gαi3 W258F mutant (**Fig. 3E**). These results demonstrate that GINIP antagonizes RGS GAP action by competing for binding to Gαi subunits.

Results presented so far from **Fig.1** to **Fig. 3** support an overarching model for the mechanism of action of GINIP on regulating GPCR signaling (**Fig. 3F**). Binding of GINIP to active Gαi subunits prevents binding of the G-protein to two other types of signaling partners: effectors of Gαi-GTP like adenylyl cyclases, and negative regulators like RGS GAPs. The latter effect prolongs the lifetime of Gαi in the GTP-bound state and dissociated from Gβγ. While in this scenario the excess of Gβγ generated is competent for signaling to effectors, Gαi-GTP is not because GINIP prevents Gαi association with adenylyl cyclase. Consequently, G-protein signaling downstream of GPCR stimulation is biased towards Gβγ over Gαi-GTP signaling by the action of GINIP (**Fig. 3F**).

### GINIP is highly expressed in brain

Next, we set out to investigate the physiological consequences of the mechanism of signaling regulation controlled by GINIP. We reasoned that, if GINIP exerts its action by modulating the response of the many GPCRs that signal via G_i_ proteins, it should have broad physiological consequences in the tissues in which it is expressed. We found that GINIP expression was restricted to the nervous system across a battery of mouse tissues, being most abundant in brain even when compared to dorsal root ganglia (DRG) (**Fig. 4A**), where GINIP has been previously shown to modulate pain processing (Gaillard et al., 2014). Motivated by this observation, we pursued the characterization of GINIP expression and function in brain. First, we found that brain-derived GINIP bound to active but not inactive Gαi3 (**Fig. 4B**), much like recombinant GINIP did (**Fig.1A**), suggesting that GINIP natively expressed in brain preserves the key function required to modulate GPCR signaling. Next, we characterized mice bearing a modified GINIP allele (i.e., GINIP 1a), which has an insertion of a *LacZ*-containing cassette in the GINIP genomic locus (**Fig. 4C**). This insertion is expected to result not only in a loss of GINIP protein (null allele), which was confirmed by western blotting (**Fig. 4D**), but also a reporter for GINIP expression. We leveraged the latter to assess the expression of GINIP in different brain regions, finding that GINIP expression is robust across the isocortex, hippocampus, striatum, amygdala, and, to a lesser extent, the thalamus (**Fig. 4E**). This expression pattern was confirmed by directly detecting GINIP expression using mRNA *in situ* hybridization in GINIP +/+ brains, for which GINIP 1a/1a served as a negative control demonstrating the specificity of the probe (**Fig. 4F**). These results suggested that GINIP may have a role in a wide range of neurons.

**Figure 4.**
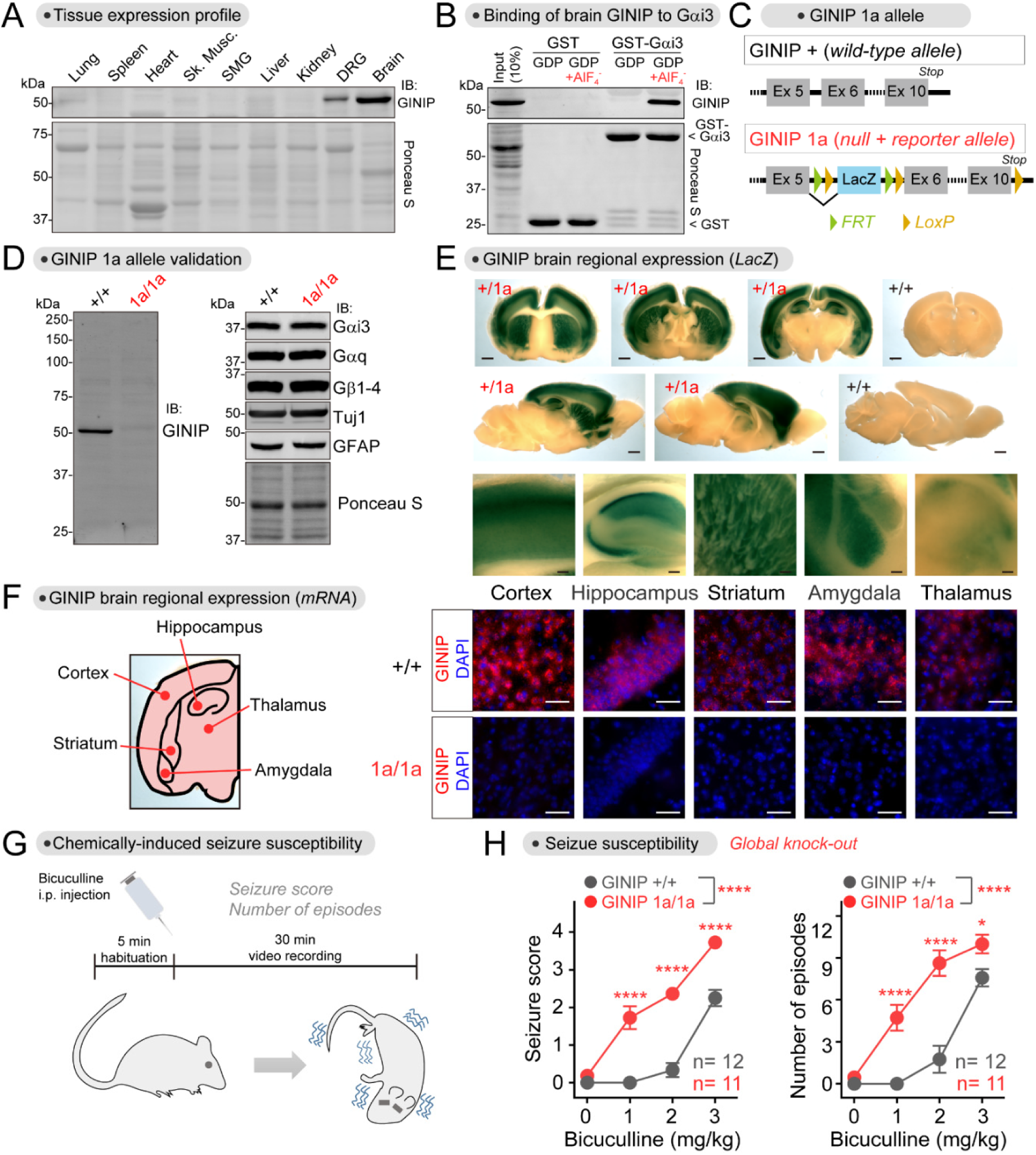
Loss of GINIP increases seizure susceptibility. **(A)** GINIP expression is restricted to the nervous system and most abundant in brain. Proteins extracted from the indicated mouse tissues were analyzed by Ponceau S staining or by immunoblotting (IB). n=2. **(B)** Active Gαi binds to brain-derived GINIP. GST or GST-Gαi3 immobilized on glutathione-agarose beads were incubated with mouse brain lysates in the presence of GDP or GDPꞏAlF_4_^−^, as indicated. Bead-bound proteins were detected by Ponceau S staining or by immunoblotting (IB). n=3. **(C)** Diagram depicting features of the GINIP 1a allele bearing a *LacZ-*containing cassette inserted between exon 5 and exon 6. **(D)** GINIP expression is ablated in GINIP 1a/1a mice without affecting the expression of several other neuronal proteins. Mouse brain lysates were analyzed by immunoblotting (IB) with the indicated antibodies. n=3. **(E)** GINIP is expressed in the cortex, hippocampus, striatum, amygdala and thalamus. β-galactosidase activity was detected by staining brain slices of GINIP +/1a or +/+ mice. Scale bars are 1 mm or 0.1 mm in images of whole brain sections or of enlarged areas, respectively. n=3. **(F)** GINIP mRNA is expressed in the cortex, hippocampus, striatum, amygdala and thalamus. GINIP mRNA was detected in mouse brain coronal slices of GINIP +/+ or 1a/1a by fluorescence *in situ* hybridization. Scale bar = 50 µm. n=3. **(G, H)** GINIP 1a/1a mice display increased susceptibility to bicuculline-induced seizures compared to GINIP +/+ mice. 10-14 week-old mice were assessed for seizure scores and number of episodes after injection of the indicated amounts of bicuculline. 11-12 mice per genotype, approximately 50/50 of males/females per group. See **Fig. S4** for results stratified by sex. Mean ± S.E.M. *p<0.05, ****p<0.0001, two-way ANOVA for genotype x concentration of bicuculline, with multiple comparisons at each concentration using Fisher’s LSD test.

### Loss of GINIP increases seizure susceptibility

Motivated by an earlier report that identified GINIP as one of the candidate genes for spontaneous, generalized seizures in a strain of epileptic rats (Kuramoto et al., 2017), we explored the susceptibility of GINIP knock-out mice to epileptic seizures. Since GINIP is highly expressed in cortical brain regions typically involved in epileptogenesis (Goldberg and Coulter, 2013), we reasoned that seizure susceptibility would be a good proxy to assess the physiological relevance of brain-expressed GINIP. Although we did not observe spontaneous seizures in GINIP 1a/1a mice compared to GINIP +/+ littermates, there were marked differences in their susceptibility to chemically-induced seizures (**Fig. 4G, H**). To induce seizures, we treated mice with bicuculline, a GABA_A_ receptor antagonist, to directly shift the balance of neurotransmission toward excitation (Chen and Toth, 2001). We found that GINIP 1a/1a mice had higher sensitivity to bicuculline-induced seizures than GINIP +/+ littermates, displaying more severe seizures and more episodes at lower concentrations of bicuculline (**Fig. 4H**). Similar differences were observed when comparing groups with mixed males and females (**Fig. 4H**) or when analyzing males and females separately (**Fig. S4A**). These findings show that GINIP is required to prevent imbalances of neurotransmission that underlie increased seizure susceptibility.

### GINIP biases GPCR-G protein signaling in neurons

We began exploring the role of GINIP in regulating neuromodulatory signaling via GPCRs in primary neurons. We cultured cortical neurons (with supportive glia) from neonatal mice and found that GINIP expression was as high as in brain tissue after 12-20 days in vitro (DIV) (**Fig. 5A**). GINIP was undetectable in cultured glia prepared in parallel from the same source (**Fig. 5B**), suggesting that GINIP is predominantly expressed in neurons. The latter was confirmed by co-staining GINIP with either a neuronal (NeuN) or a glial (GFAP) marker. We found that almost all NeuN^+^ cells expressed GINIP, whereas no GFAP^+^ cell expressed GINIP (**Fig. 5C**). To investigate the role of GINIP in GPCR signaling in neurons, we generated mice bearing a floxed GINIP allele (GINIP flox) (**Fig. S5A-C**). Cortical neurons cultured from GINIP flox/flox mice were transduced with an adeno-associated virus (AAV) expressing Cre recombinase (AAV-Cre) to generate GINIP null cells that were compared to uninfected control cells (**Fig. S5D-E**). To monitor GPCR responses, we expressed a BRET biosensor named BERKY that is suitable for the detection of endogenous free Gβγ in response to endogenous neurotransmitter receptors in neurons (Maziarz et al., 2020). We found that cells lacking GINIP had reduced Gβγ responses upon stimulation of two different G_i_-coupled GPCRs, GABA_B_R or α2-AR, with their respective selective agonists (**Fig. 5D-E**). In contrast, Gβγ responses elicited by stimulation with an agonist for β-adrenergic receptors, which couple to G_s_ instead of G_i_ proteins, were unaffected by the loss of GINIP under the same experimental conditions (**Fig. 5F**). These results demonstrate that GINIP endogenously expressed in neurons promotes Gβγ signaling in response to GPCR stimulation, much like in cell lines expressing exogenous GINIP (**Fig. 3**). Next, we investigated if GINIP would also modulate cAMP changes in response to stimulation of a G_i_-activating GPCR, like previously found in cell lines (**Fig. 2**). For this, we expressed a BRET-based biosensor for cAMP (Masuho et al., 2015) in neurons of GINIP flox/flox mice and compared responses with and without Cre-mediated ablation of GINIP as done in the free Gβγ measurements above. We found that the inhibition of forskolin-stimulated cAMP after stimulation of GABA_B_R was enhanced upon loss of GINIP (**Fig. 5G**). These results suggest that, much like in cell lines, GINIP dampens Gαi-GTP-mediated inhibition of cAMP production by adenylyl cyclase. Collectively, these observations indicate that GINIP biases inhibitory GPCR responses in neurons by favoring Gβγ over Gαi signaling.

**Figure 5.**
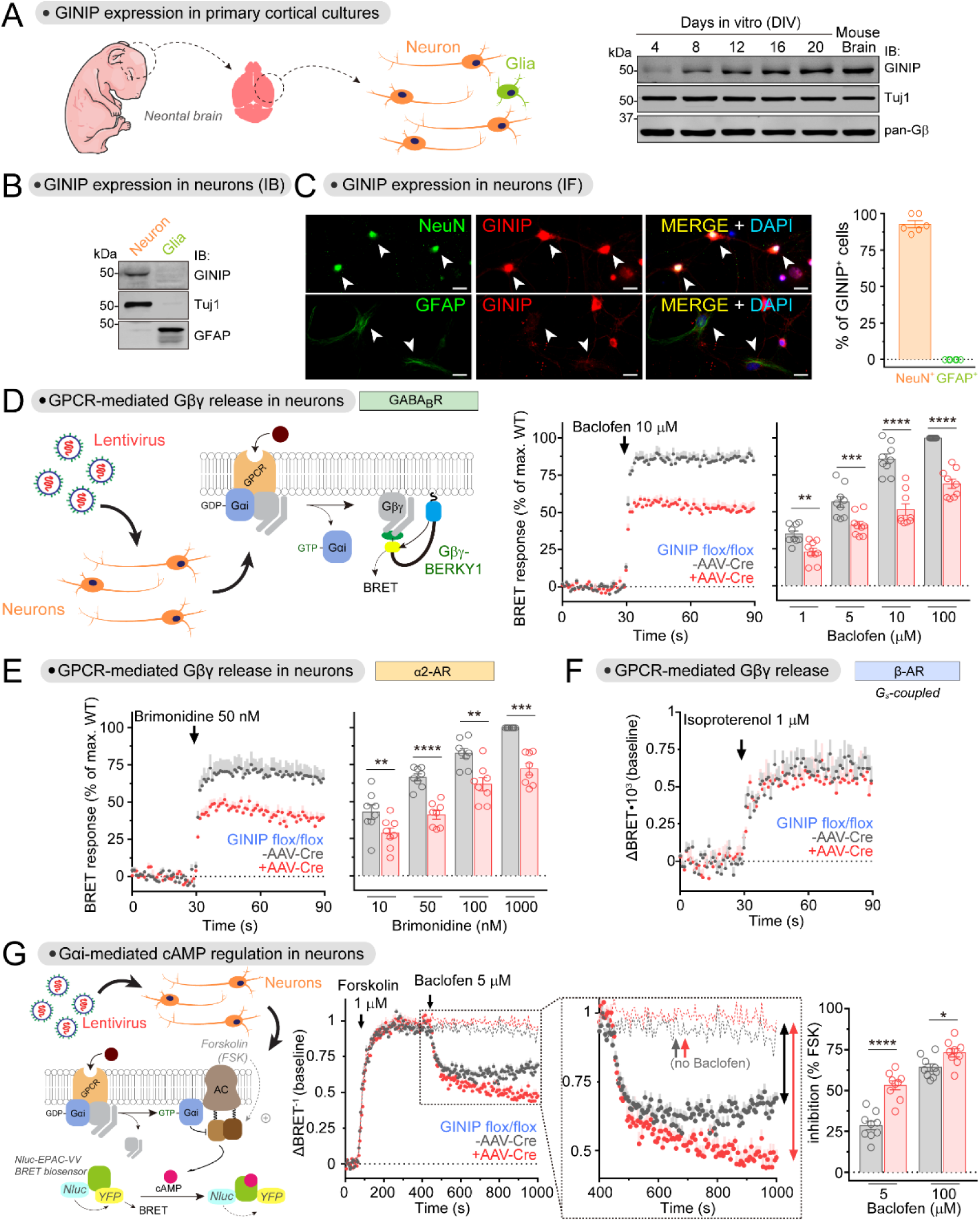
GINIP regulates G_i_-coupled GPCR signaling in neurons. **(A)** GINIP is expressed in cortical neuron cultures. Neuron cultures established from the cortices of neonatal mouse brains were analyzed by immunoblotting (IB) on the indicated days in vitro (DIV). n=2. **(B)** GINIP is not expressed in cortical glial cultures. Neuron and glial cultures established from the cortices of neonatal mouse brains were analyzed by immunoblotting (IB) on DIV12. n=3. **(C)** GINIP is expressed in cortical neurons but not in glia. GINIP was co-stained with NeuN or GFAP in DIV12 cortical cultures. Arrows indicate NeuN^+^ or GFAP^+^ cells. The proportion of in NeuN^+^ or GFAP^+^ cells that were GINIP^+^ was quantified from three independent cultures (two image fields per experiment). Mean ± S.E.M. Scale bar = 20 µm. **(D, E)** Loss of GINIP decreases Gβγ responses triggered by GABA_B_R (D) and α2-AR (E). BRET was measured in DIV12-14 cortical neurons from GINIP flox/flox mice that had been infected (red) or not (black) with an AAV-Cre virus. Kinetic traces of one concentration of agonist are shown on the left of each panel, and the quantification of BRET response amplitudes at different concentrations of agonist is shown on the right. BRET responses were normalized to the maximum response of WT in each experiment. Mean ± S.E.M., n=8-9. **p<0.01, ***p<0.001, ****p<0.0001, paired t-test. **(F)** Loss of GINIP does not affect Gβγ responses triggered by β-AR. BRET was measured in DIV12-14 cortical neurons from GINIP flox/flox mice that had been infected (red) or not (black) with an AAV-Cre virus. Mean ± S.E.M. n=5 **(G)** Loss of GINIP enhances the inhibition of adenylyl cyclase upon stimulation of GABA_B_R. cAMP was measured by BRET in DIV16-18 cortical neurons from GINIP flox/flox mice that had been infected (red) or not (black) with an AAV-Cre virus. Kinetic traces of one concentration of baclofen are shown on the left, and the quantification of inhibition of forskolin (FSK)-induced cAMP at different concentrations of baclofen is shown on the right. Dotted lines in the kinetic traces indicate controls not stimulated with baclofen. Mean ± S.E.M., n=9. *p<0.05, ****p<0.0001, paired t-test.

### GINIP localizes to inhibitory but not excitatory synapses

Close inspection of cortical neurons in culture immunostained for GINIP revealed that the protein localized in puncta that partially overlapped with the synaptic marker synaptophysin (SYP) (**Fig. 6A**). The immunostaining was specific for GINIP because no signal was detected in neurons from GINIP 1a/1a mice (**Fig. 6A**). Further experiments co-staining with other markers revealed that GINIP is expressed in both inhibitory and excitatory neurons, as determined by the expression of the GABAergic marker GAD65 or the glutamatergic marker vGlut1, respectively (**Fig. 6B**). However, GINIP colocalizes with markers of inhibitory but not excitatory synapses (**Fig. 6C**). For example, ~60-80% of GINIP^+^ puncta in dendrites colocalized with GAD65, a marker of inhibitory presynaptic terminals, and ~30% colocalized with gephyrin, a marker of inhibitory postsynaptic structures (**Fig. 6C**). In contrast, GINIP did not colocalize with vGlut1 or PSD95, which mark excitatory presynaptic and postsynaptic structures, respectively (**Fig. 6C**). Interestingly, the co-localization of SYP was bimodal— i.e., in dendrites of some neurons ~70% of GINIP^+^ puncta were positive for SYP, whereas co-localization was absent in dendrites of other neurons (**Fig. 6C**). In light of results obtained with other markers and the fact that SYP marks both inhibitory and excitatory presynaptic terminals, the most likely explanation for this is that GINIP co-localizes with SYP at inhibitory presynaptic terminals (like those with GAD65) but not at excitatory presynapses (like those with vGlut1). In summary, our results indicate that, even though GINIP is expressed in both excitatory and inhibitory cortical neurons, its subcellular distribution is restricted to inhibitory synapses, probably at both presynaptic and postsynaptic structures.

**Figure 6.**
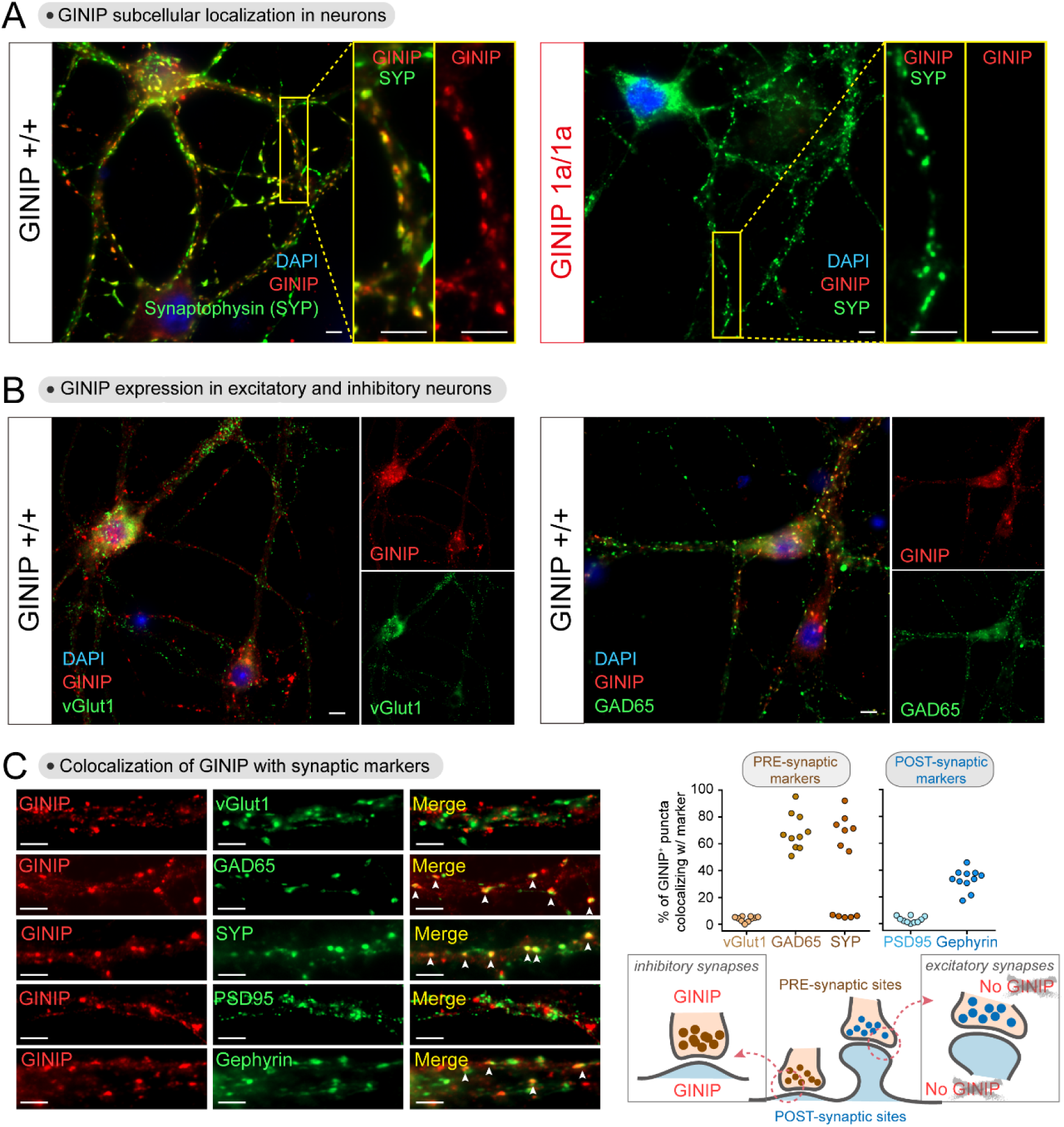
GINIP localizes to inhibitory but not excitatory synapses. **(A)** GINIP protein localizes to dendritic puncta in cortical neurons. DIV21 cortical neurons from GINIP +/+ or 1a/1a mice were co-stained for GINIP and synaptophysin (SYP) before fluorescence imaging. Yellow boxes indicate areas enlarged on the right side of the main images. Enlarged views of representative neurites (yellow box) are shown on the right side of the main images. **(B)** GINIP is expressed in both excitatory (vGlut1^+^) and inhibitory (GAD65^+^) neurons. DIV21 cortical neurons from GINIP +/+ mice were co-stained for GINIP and the indicated markers before fluorescence imaging. **(C)** GINIP co-localizes with markers of inhibitory but not excitatory synapses. *Left*, cortical neurons co-stained for GINIP and the indicated markers of different synaptic compartments were imaged by confocal fluorescence microscopy. *Top right*, quantification of the colocalization of GINIP with synaptic markers. Scatter plot values are the percentage of GINIP positive puncta that were positive for each synaptic marker of one field. Data correspond to 3-4 fields from 3 independent experiments. *Bottom right*, diagram depicting the proposed localization of GINIP in different synaptic sites. All scale bars are 10 µm. All results presented are representative of n ≥ 3 experiments.

### GINIP affects GPCR-mediated neuromodulation in both excitatory and inhibitory neurons

Since GINIP is expressed in both excitatory and inhibitory cortical neurons in culture (**Fig. 6**), next we asked if GINIP regulated GPCR neuromodulation in excitatory or inhibitory neurons, or in both. For this, we set out to carry out patch-clamp electrophysiological recordings in brain slices. We first confirmed that GINIP is expressed in both excitatory and inhibitory neurons of the cortex (**Fig. 7A**) by using mRNA *in situ* hybridization with brain slices. Next, we evaluated the impact of GINIP loss on neuromodulatory influence of GABA_B_R using patch-clamp recordings of excitatory cortical pyramidal neurons in brain slices. Activation of postsynaptic GABA_B_R in these neurons reduces excitability by activating potassium current thought to be largely mediated by G protein-gated inwardly rectifying K^+^ (GIRK) channels, a canonical Gβγ effector (Betke et al., 2012; Luscher and Slesinger, 2010). Indeed, application of GABA_B_R agonist baclofen produced prominent K^+^ current (**Fig. 7B**). The amplitude of this current was significantly reduced in GINIP 1a/1a compared to GINIP +/+ suggesting loss in the efficiency of postsynaptic GIRK channel activation by Gβγ in the absence of GINIP (**Fig. 7B**).

**Figure 7.**
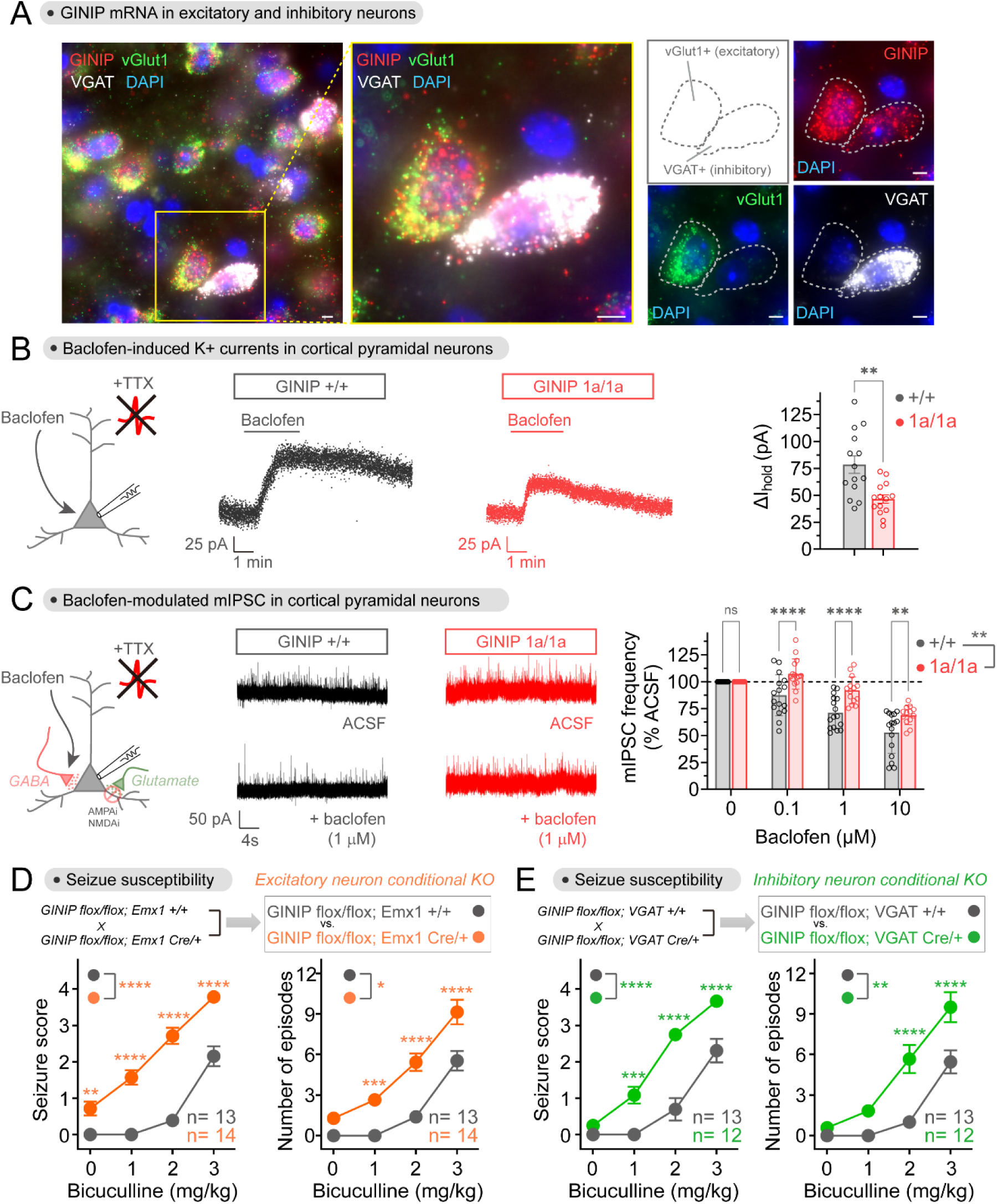
Loss of GINIP from either excitatory or inhibitory neurons affects inhibitory neuromodulation and increases seizure susceptibility. **(A)** GINIP mRNA is expressed in excitatory and inhibitory cortical neurons. GINIP, vGlut1 and VGAT mRNAs were simultaneously detected in mouse cortical slices by fluorescence in situ hybridization. All scale bars are 10 µm. Results presented are representative of n ≥ 3 experiments. **(B)** Loss of GINIP reduces GIRK currents in response to baclofen. Representative traces of baclofen-induced holding current change in cortical pyramidal neurons from GINIP +/+ (black) or GINIP 1a/1a (red) slices are shown in the left and middle, whereas quantification of peak amplitude across multiple cells is shown on the right. Mean ± S.E.M. (n=14 per group), **p<0.01, unpaired t-test. Baclofen = 50 μM. **(C)** Loss of GINIP dampens baclofen-induced reduction of mIPSC frequency. Representative traces of mIPSC recorded from GINIP +/+ (black) and GINIP 1a/1a (red) cortical pyramidal neuron before and after baclofen are shown in the left and middle, whereas quantification of mIPSC frequency for different concentrations of baclofen relative to controls is shown on the right. Mean ± S.E.M. n=13-16 per group. **p<0.01, ***p<0.001, ****p<0.0001, two-way ANOVA for GINIP genotype x baclofen concentration, with multiple comparisons at each concentration using Fisher’s LSD test. **(D, E)** Loss of GINIP from Emx1^+^ (excitatory) neurons (B) or from VGAT^+^ (inhibitory) neurons (C) results in increased seizure susceptibility. GINIP flox/flox mice were compared to littermates bearing Emx1 Cre or VGAT Cre driver alleles to achieve neuron-specific ablation of GINIP. 10-14 week-old mice were assessed for seizure scores and number of episodes after injection of the indicated amounts of bicuculline. 12-14 mice per genotype, approximately 50/50 of males/females per group. See **Fig. S4** for results stratified by sex. Mean ± S.E.M. *p<0.05, **p<0.01, ***p<0.001, ****p<0.0001, two-way ANOVA for genotype x concentration of bicuculline, with multiple comparisons at each concentration using Fisher’s LSD test.

To evaluate the effect of GINIP in the inhibitory neurons, we measured inhibition of GABA release mediated by activation of presynaptic GABA_B_R autoreceptors. Here, Gβγ released upon activation of GABA_B_R inhibits neurotransmission by inhibiting Ca^2+^ ion channels and by blocking SNARE-mediated vesicular fusion (Betke et al., 2012; Zurawski et al., 2019). For this, we recorded miniature inhibitory postsynaptic currents (mIPSC) in pyramidal cortical neurons in the presence or absence of baclofen, reflecting neurotransmitter release from inhibitory, GABAergic neurons. As expected, application of baclofen led to a reduction of the frequency of mIPSCs in GINIP +/+ (**Fig. 7C**). However, in GINIP 1a/1a, this baclofen-induced reduction of mIPSC frequency was significantly lower than in GINIP +/+ (**Fig. 7C**). We found no differences by genotype in baseline mIPSC frequencies without baclofen treatment (4.72 Hz versus 4.75 Hz in GINIP 1a/1a and GINIP +/+, respectively, n=13-15 per group, p>0.05 unpaired t-test). Taken together, these results suggest that GINIP facilitates GPCR-mediated modulation of inhibitory neurotransmission in both excitatory and inhibitory neurons.

### Loss of GINIP from either excitatory or inhibitory neurons increases seizure susceptibility

Based on the observed effects of GINIP on regulating neuromodulatory responses in both excitatory and inhibitory neurons, next we investigated if the phenotype of higher seizure susceptibility observed in global GINIP knock-out mice (**Fig. 4G, H**) was due to the function of GINIP in either excitatory or inhibitory neurons, or in both. For this, we crossed GINIP flox mice with different Cre driver lines to specifically ablate GINIP expression in excitatory or inhibitory neurons across the brain regions where GINIP is expressed. For the former, crosses were made with Emx1 Cre mice (Gorski et al., 2002), whereas VGAT Cre line (Vong et al., 2011) was used for the latter. We confirmed specific loss of GINIP from excitatory (vGlut1^+^) or inhibitory neurons (VGAT^+^) in GINIP flox/flox mice bearing the Emx1 Cre or VGAT Cre allele, respectively, across all brain regions investigated (**Fig. S6**). We found that loss of GINIP from either excitatory or inhibitory neurons had a similar effect on bicuculine-induced seizure susceptibility, and that this effect was also similar to that observed with global GINIP knock-out (**Fig. 4G, H**)— i.e., it caused more severe seizures and more episodes at lower concentrations of bicuculline (**Fig. 7E-F**). As for the global GINIP knock-out, the increase in seizure susceptibility was observed when comparing groups with mixed males and females (**Fig. 7E-F**) or when analyzing males and females separately (**Fig. S4B-C**). These findings demonstrate that GINIP is required both in excitatory and inhibitory neurons to prevent imbalances of neurotransmission that underlie increased seizure susceptibility.

## DISCUSSION

The main advance provided by this work is the identification of a unique mechanism of G-protein regulation that sets the tone of neurotransmission by fine tuning inhibitory neuromodulation triggered by GPCRs. This mechanism is mediated by the Gαi-binding protein GINIP, but fundamentally differs from other regulators that bind Gα subunits in that GINIP does not affect directly the enzymatic activity of the G-protein. Instead, when GINIP binds tightly to a region of active Gαi comprising the α3 helix and the SwII (**Fig. 1**), it competes simultaneously with the binding of effectors (**Fig. 2**) and of RGS GAPs (**Fig. 3**). This leads to a GPCR signaling outcome that is apparently paradoxical because some G-protein signals are enhanced while others are inhibited— i.e., G_i_ responses are biased to enhance the effects one of its active signaling species (free Gβγ) in detriment of the effects of the other active signaling species (Gαi-GTP). This is explained by the prolonged lifetime of Gαi in its GTP-bound form upon RGS GAP displacement by GINIP, which is accompanied by the corresponding dissociation of Gβγ. However, Gαi-GTP in this scenario remains “sequestered” by GINIP in a complex that is incompetent for engagement with its effector adenylyl cyclase. We show that this mechanism has a broad effect on GPCR signaling that is not limited to the previously described regulation GABA_B_R signaling by GINIP in the peripheral nervous system (Gaillard et al., 2014), but that potentially encompasses the regulation of any G_i_-coupled GPCR co-expressed with GINIP across different brain regions (**Fig. 4**). In this regard, based on its broad expression across different major types of neurons, like GABAergic and glutamatergic neurons (**Fig. 6, Fig. 7**), GINIP is poised to exert circuit level effects on neurotransmission in brain by controlling GPCR-mediated neuromodulation, an idea supported by our signaling, electrophysiological, and behavioral results (**Fig. 5**, **Fig. 7**).

The mechanism by which GINIP regulates GPCR signaling is different from those described for other regulators that bind to Gα subunits, like GDIs or GAPs. A first distinctive feature is that GINIP does not affect directly the ability of Gα subunits to bind and/ or hydrolyze nucleotides. Another distinctive feature is that GINIP has effects of opposite sign on signaling branches triggered by the same receptor depending on whether they are mediated by Gα or Gβγ subunits, whereas other regulators do not discriminate GPCR responses this way. For example, a previous report showed that the GDI LGN could enhance basal GIRK channel activation in neurons via Gβγ subunits released upon its binding to inactive, GDP-bound Gα in the absence of GPCR stimulation, but that GIRK responses were dampened by LGN upon GPCR stimulation (Wiser et al., 2006). Together, these observations suggest that GDIs like LGN inhibit all G-protein responses triggered by a GPCR by disrupting G-protein heterotrimers, which are the obligatory substrate for GPCRs. RGS GAPs also work as negative regulators of GPCR signaling regardless of whether the readout depends on Gα-GTP or Gβγ (Masuho et al., 2020).

Much attention has been paid in the recent years to the concept of ligand-induced bias in GPCR signaling (Kenakin, 2019; Kolb et al., 2022). The underlying idea is that the strength of different responses triggered by a GPCR can be modulated to different extents depending on the properties of the ligand that activates the receptor. One general hypothesis is that different ligands can facilitate different GPCR conformations, which in turn are better suited to engage preferentially some transducers over others. Most work has focused on differences between engagement of G-proteins vs β-arrestins, although there could also be ligand-induced bias for different types of G-proteins. Here, we describe a different type of signaling bias. Instead of regulating the transducer that engages the GPCR, GINIP determines the functional outcome by biasing signaling mediated by different G-protein subunits. This is not necessarily determined by the nature of the ligand. Instead, the work presented here is a case of “system bias”— i.e., the relative sensitivity of pathways activated by the receptor that are hard-wired by the physiology of the system (Kenakin, 2018, 2019). Although it is becoming recognized that system bias is a general impediment for the translation of *in vitro* pharmacology into useful therapeutics *in vivo* (Kenakin, 2018), little is known about the mechanisms underlying this class of bias beyond attributing it to differences in receptor and transducer stoichiometry. Recent work has put forth the idea that receptor proximal events, like regulation of G-proteins by RGS GAPs, could also contribute to system bias by shaping the relative strength of GPCR signals mediated by different types of G-proteins in neurons or cardiac cells (Anderson et al., 2020; Masuho et al., 2020). Our work presented here provides a detailed molecular understanding of a mechanism of system bias that operates specifically in neurons through the regulation of receptor proximal events to determine the sensitivity of different pathways under the control of not only the same receptor, but also the same G-protein type. A question that remains open is how the action of GINIP could be modulated to tune this bias. One obvious mechanism is through control of GINIP expression in specific neuron populations of different brain regions, which remains to be characterized in detail. However, even when expressed in a particular type of neuron, GINIP localization to different subcellular compartments (like synapses, **Fig. 6**) will also determine the specific contexts in which GPCR signaling bias occurs.

The increased susceptibility to bicuculline-induced seizures observed upon loss of GINIP suggests an overall decrease in neuroinhibition, which is in agreement with other results from electrophysiological and G-protein signaling experiments in neurons (**Fig. 5**, **Fig. 7**). Our behavioral results are consistent with and expand on previous observations in male rats deficient for GINIP expression, which displayed increased spontaneous seizures (Kuramoto et al., 2017) or seizures induced using an experimental paradigm different from ours (Serikawa et al., 2019). We not only observed similar effects in mice of both sexes, but also were able to leverage a conditional GINIP floxed allele to learn that loss of GINIP from either excitatory or inhibitory neurons increases similarly seizure susceptibility (**Fig. 7**). While this indicates an important role for GINIP in regulating the overall balance of neurotransmission when acting in any of these two broad neuron types, the mechanisms leading to a similar systems-level phenotype of decreased neuroinhibition are unclear. One possibility is that non-cell autonomous, circuit-level mechanisms lead to the similar overall phenotype. However, we cannot rule out that cell-autonomous Gβγ vs Gαi-GTP responses regulated by GINIP are integrated differently depending on the type of neuron. Since both Gβγ and Gαi-GTP signaling tend to be neuroinhibitory but are regulated in opposite directions by GINIP, the net effects on regulating neurotransmission would depend on how they are integrated in a particular cell type.

Gαo (in particular the GαoA isoform) is the most abundant G-protein in brain, and GPCRs that activate G_i_ almost invariably also activate G_o_ (Hauser et al., 2022). Yet, GINIP regulates responses by the former but not the latter. Why have a mechanism in place to regulate two closely related G-protein types that tend to be co-activated? While both GαoA and Gαi isoforms give rise to Gβγ subunits that might be functionally similar upon stimulation of same GPCRs, the key functional difference of GαoA with respect to Gαi is that it does not regulate adenylyl cyclase directly. It is tempting to speculate that GINIP comes into play to regulate GPCR responses by the G-protein subtype for which coordination between Gβγ and Gα-GTP is relevant— i.e., G_i_. In this regard, it is also important to note that while GPCRs that activate G_i_ also co-activate G_o_, and that both G-proteins contribute to the formation of Gβγ, regulation of G_i_ by GINIP affects the overall levels of free Gβγ (**Fig. 5**) or Gβγ-dependent responses like post-synaptic GIRK channel regulation or pre-synaptic regulation of neurotransmitter release (**Fig. 7**). This suggests that despite the abundance of GαoA in neurons, a significant fraction of Gβγ-dependent signaling is mediated through Gαi isoforms in brain.

In summary, this work provides detailed mechanistic insights into how GPCR-mediated neuromodulation is fine-tuned by coordinating the level of responses triggered by different G-protein subunits. By understanding the molecular basis of how receptor-mediated signaling is hard-wired in the native context of neurons, we might be able to envision better ways to pharmacologically target neuroaalogical and neuropsychiatric disorders.

## ACKNOWLEDGEMENTS

This work was primarily supported by NIH grant R01NS117101 (to MG-M). AL is supported a F31 Ruth L. Kirschstein NRSA Predoctoral Fellowship (F31NS115318). We thank Vickery Trinkaus-Randall (Boston University) for access to confocal microscopes. We thank the following investigators for providing DNA plasmids: N. Lambert (Augusta University, Augusta, GA), J. Blumer (Medical University of South Carolina, Charleston, SC), J. Sondek (University of North Carolina, Chapel Hill, NC), and M. Linder (Cornell University). We also thank C. Dessauer (University of Texas Health Science Center at Houston, TX) for providing plasmids and protocols for the in vitro adenylyl cyclase experiments. We thank members of the Garcia-Marcos laboratory for reading a draft of this manuscript and making comments.

## Author Contributions

J-C.P., A.L., M.D., A.S., M.P.P., S.B. and M.G-M. conducted experiments. J-C.P., A.L., M.D., A.S., H.Y., K.A.M. and M.G-M designed experiments and analyzed data. J-C.P., A.L., and M.G-M. wrote the manuscript with input from all authors. M.G-M. conceived and supervised the project.

## Data and materials availability

All data are available in the main text or the supplementary materials.

## Methods

### Plasmids

Plasmids for the bacterial expression of His-tagged human GINIP (pLIC-His-GINIP) or RGS4 (pLIC-His-RGS4) were generated using a previously described ligation-independent cloning (LIC) system (Stols et al., 2002). Briefly, insert sequences containing LIC compatible flanking regions were amplified by PCR and inserted into the pLIC-His plasmid generously provided by D. Siderovski (UNT Health Science Center). Similarly, an LIC-compatible GINIP amplicon was inserted into the pLIC-GST plasmid kindly provided by J. Sondek (University of North Carolina at Chapel Hill) (Cabrita et al., 2006) to generate pLIC-GST-GINIP. Plasmids for the bacterial expression of His-Gαi3 (rat), GST-Gαi3, His-Gαi1, His-Gαi2, and His-Gαo have been described previously (Garcia-Marcos et al., 2009; Garcia-Marcos et al., 2020; Marivin et al., 2020). The plasmid pET24a-Gαi3 used for the bacterial expression of human His-Gαi3 was a kind gift of I. Shimada (de Opakua et al., 2017). A plasmid encoding rat Gαi1 containing a hexahistidine tag in the b/c loop (Gαi1-6xHis(int), required for producing myrisotoylated Gαi1) was kindly provided by C. Dessauer (UT Southwestern) (Dessauer et al., 1998). Gαi1-6xHis(int) was subcloned without additional affinity tags between the NdeI and BglII sites of the pLIC-His vector (pLIC-Gαi1-6xHis(int)). The pbb131 plasmid encoding yeast *N*-myristoyltransferase (NMT) (Mumby and Linder, 1994) was a gift from Maurine Linder (Cornell University). A plasmid encoding the C1 domain (residues 444-751) of canine adenylyl cyclase 5 (AC5) was provided by C. Dessauer (UT Southwestern) (Dessauer et al., 1998). AC5 C1 (with a C-terminal His tag) was subcloned between the NcoI and XhoI sites of the bacterial expression vector pET28b to generate pET28b-AC5 C1 without adding any other affinity tag. pET15b-hAC2 C2 encoding residues 871-1082 of human adenylyl cyclase 2 (AC2 C2) with an N-terminal His tag was provided by K. Shokat (University of Californa San Francisco) (Hu and Shokat, 2018). Plasmids for bacterial expression of GST-GAIP and GST-KB1753 have been described previously (Garcia-Marcos et al., 2010; Leyme et al., 2017). The plasmid for the expression of a His–tagged short isoform of bovine Gαs (pHis6-Gαs) in bacteria was kindly provided by N. Artemyev (University of Iowa). Plasmids for expression of GST-Gαi3/o chimeras have been described previously (Garcia-Marcos et al., 2010). The plasmid for the expression of His-DAPLE CT, pET28b-DAPLE (1650-2028), was described previously (Aznar et al., 2015).

The plasmid for mammalian expression of C-terminally 3xFLAG-tagged GINIP (GINIP-FLAG) was generated for this paper by inserting the human GINIP sequence between EcoRI and BamHI sites of the p3xFLAG-CMV-14 vector and replacing the stop codon by a serine (p3xFLAG-CMV-14-GINIP). Plasmids encoding for C-terminally 3xFLAG-tagged Gαi3, untagged Gαi3, Gαo, and Gαs were described previously (Beas et al., 2012; Garcia-Marcos et al., 2010; Garcia-Marcos et al., 2011b; Ghosh et al., 2008), whereas the plasmid for the expression of EE-tagged Gαz was purchased from the cDNA Resource Center (GNA0Z0EI00) (Bloomsberg University, PA). The plasmid for mammalian expression of Gαq internally tagged with HA (pcDNA3-Gαq-HA) was kindly provided by P. Wedegaertner (Thomas Jefferson University) (Wedegaertner et al., 1993), whereas the plasmid for mammalian expression of Gα12 internally tagged with MYC (pcDNA3.1-Gα12-MYC) was from T. Meigs (UNC Asheville) (Meigs et al., 2005). The plasmid for mammalian expression of YFP-tagged human adenylyl cyclase 5 (pcDNA3-YFP-hAC5) was a gift from C. Dessauer (UT Southwestern) (Sadana and Dessauer, 2009). The plasmid for mammalian expression of the long isoform of the human Dopamine 2 receptor (pcDNA3.1(+)-FLAG-D2DR) was provided by A. Kovoor (University of Rhode Island). The plasmid encoding α2_A_-AR (pcDNA3-α2_A_-AR) has been described previously (Oner et al., 2010). pcDNA3.1(+)-GABA_B_R1a and pcDNA3.1(+)-GABA_B_R2 were a gift from Paul Slessinger, Mount Sinai NY. The following mammalian expression plasmids were described by us previously: pcDNA3.1-hRGS7, pcDNA3.1-Gβ5, pcDNA3.1-R7BP, pcDNA3.1-masGRK3ct-Nluc, pcDNA3.1-Nluc-EPAC-VV (Masuho et al., 2015). pcDNA3.1-Venus(1–155)-Gγ_2_ (VN-Gγ2), and pcDNA3.1-Venus(155-239)-Gβ1 (VC-Gβ1) were a gift from N. Lambert (Augusta University, GA) (Hollins et al., 2009). pcDNA3.1(-)-3xHA-RGS8 was acquired from the cDNA Resource Center (RGS080TN00) (Bloomsberg University, PA). pcDNA3-GAIP was a gift from M. Farquhar (De Vries et al., 1996). A plasmid encoding rat Gαi3 tagged with Nluc in the a/b loop, pcDNA3.1(-)-Gαi3-Nluc(a/b), was generated by inserting EcoRI and XhoI restriction sites between residues 91 and 92 of Gαi3, and then inserting Nluc by Gibson assembly at those sites, which were maintained in the final construct. The resulting construct contains a 5’ linker EF and 3’ linker SS flanking the Nluc sequence.

Lentiviral packaging plasmids psPAX2 and pMD2.G, and the plasmid to produce a lentivirus for the expression of the Gβγ-BERKY1 biosensor in neurons (pLenti-hSyn-Gβγ-BERKY1) have been described previously (Maziarz et al., 2020). The plasmid to produce lentiviral particles for the expression of the cAMP BRET biosensor Nluc-EPAC-VV in neurons was generated by amplifying the sequence from pcDNA3.1-Nluc-EPAC-VV and inserting it between the AgeI and EcoRI sites of pLenti-hSynapsin-Cre-WPRE (Addgene #86641;(Sakurai et al., 2016)) by Gibson assembly.

### Protein expression and purification

His-tagged and GST-tagged proteins were expressed in BL21(DE3) *E. coli* transformed with the corresponding plasmids by overnight induction at 23 °C with 1 mM isopropyl β-D-1-thio-galactopyranoside (IPTG), with the exception of His-GINIP and GST-GINIP, which were induced for 5 hours at 23 °C. IPTG was added when the OD_600_ reached ~0.8. Unless otherwise indicated, protein purification was carried out following previously described protocols (Aznar et al., 2015; Garcia-Marcos et al., 2009). Briefly, bacteria pelleted from 1 liter of culture were resuspended at 4 °C in 25 ml of lysis buffer (50 mM NaH_2_PO_4_, pH 7.4, 300 mM NaCl, 10 mM imidazole, 1% (v/v) Triton X-100, supplemented with a protease inhibitor mixture of 1 μM leupeptin, 2.5 μM pepstatin, 0.2 μM aprotinin, and 1 mM phenylmethylsulfonyl fluoride). When purifying Gα subunits, this buffer was supplemented with 25 μM GDP and 5 mM MgCl_2_. After sonication (4 pulses of 30 s separated by 30 s intervals for cooling), the lysate was cleared by centrifugation at 12,000 × *g* for 30 min at 4 °C. The soluble fraction was used for affinity purification in batch on HisPur Cobalt (Thermo, 89964) or GSH-agarose resins (Thermo, 16100) by incubating lysate and beads with rotation for 2 hours at 4 °C. Resin was washed 3 times with lysis buffer and then eluted with lysis buffer supplemented with 250 mM imidazole or with 50 mM Tris-HCl, pH 8, 100 mM NaCl, 30 mM reduced GSH, respectively. Proteins were buffer exchanged to PBS (137 mM NaCl, 2.7 mM KCl, 8 mM Na_2_HPO_4_, and 2 mM KH_2_PO_4_) by overnight dialysis (10,000 Da cut-off) at 4 °C, except for Gα proteins, which were buffer exchanged to 20 mM Tris-HCl, pH 7.4, 20 mM NaCl, 1 mM MgCl_2_, 1 mM DTT, 10 μM GDP, 5% (v/v) glycerol using a HiTrap desalting column (Cytiva, 29048684) connected to an AKTA FPLC. All protein samples were aliquoted and stored at −80 °C. Gαi1^RM/AS^ was stored as single use aliquots for experiments to avoid freeze/thaw cycles.

Myristoylated Gαi1 (myr-Gαi1) was purified as described above from BL21(DE3) *E. coli* bacteria co-expressing the plasmid encoding Gαi1-6xHis(int) with a plasmid encoding N-myristoyl transferase (NMT), except that after the cobalt affinity purification step the eluate was subjected to ion-exchange chromatography in a HiTrapQ HP column (Cytiva, 17115401). AC5 C1 was induced and purified as described above for other His-tagged proteins except that 1 mM β-mercaptoethanol was present in all purification buffers and after the cobalt affinity purification step the eluate was subjected to ion-exchange chromatography in a HiTrapQ HP column. AC2 C2 was expressed by overnight induction at 23 °C with 40 μM IPTG. Protein was purified as described above for other His-tagged proteins except that after the cobalt affinity purification step the eluate was subjected to ion-exchange chromatography in a HiTrapQ HP column. AC5 C1 and AC2 C2 were then buffer exchanged to 20 mM HEPES pH 8.0, 1 mM EDTA, 20 mM NaCl, 2 mM DTT, and 5% (v/v) glycerol in a HiTrap desalting column connected to an AKTA FPLC and aliquoted before storage at −80 °C.

Purification of His-Gαs was carried out using a previously described protocol (Leyme et al., 2014b) that differs from the one described above for other His-tagged proteins. Briefly, His-Gαs was expressed in BL21(DE3) *E. coli* transformed with the corresponding plasmid by overnight induction at 23 °C with 0.1 mM IPTG when OD_600_ reached ~0.5. Bacteria were pelleted and resuspended at 4 °C in lysis buffer (50 mM Tris-HCl, pH 8.0, 50 mM NaCl, 5 mM MgCl_2_, 50 μM GDP, and 5 mM β-mercaptoethanol, supplemented with a protease inhibitor mixture of 1 μM leupeptin, 2.5 μM pepstatin, 0.2 μM aprotinin, and 1 mM PMSF). After sonication, the lysate was cleared by centrifugation at 12,000 × *g* for 30 min at 4 °C. The supernatant was adjusted to 500 mM NaCl and 20 mM imidazole before affinity purification by incubation with nickel-nitrilotriacetic acid (Ni-NTA) resin (Qiagen, 30210) for 90 min at 4 °C. Resin was washed four times with lysis buffer, and protein was eluted with lysis buffer supplemented with 100 mM imidazole. The eluted fraction was adjusted to 50 mM Tris-HCl, pH 8.0, 50 mM NaCl, 5 mM MgCl_2_ using a protein concentrator with a 10-kDa cutoff (Millipore, UFC801024 before loading onto a HiTrap Q HP column connected to an ÄKTA FPLC. Proteins were eluted by applying a 50–500 mM NaCl gradient, and fractions containing His-Gαs were pooled and supplemented with 10 μM GDP and 5 mM β-mercaptoethanol before concentration in 50 mM Tris-HCl, pH 8.0, 150 mM NaCl, 5 mM MgCl_2_, 5% (w/v) glycerol, 10 μM GDP, and 5 mM β-mercaptoethanol, and storage at −80 °C.

### Pulldown assays

The following GST-fused proteins were immobilized on GSH-agarose beads (Thermo, 16100) for 90 min at room temperature in PBS (range of protein amounts used in different experiments is in parenthesis): GST (2-20 μg), GST-GINIP (4-12 μg), GST-Gαi3 (1-8 μg), GST-KB1753 (15 μg), GST-GAIP (3 μg). Beads were washed twice with PBS and resuspended in 300-400 μl of binding buffer (50 mM Tris-HCl, pH 7.4, 100 mM NaCl, 0.4% (v/v) Nonidet P-40, 5 mM EDTA, 2 mM DTT) supplemented with the following additives depending on the conditions indicated in figures and legends: 30 μM GDP (GDP condition), or 30 μM GDP, 30 μM AlCl_3_, and 10 mM NaF (GDPꞏAlF_4_^−^ condition), or 30 μM GTPγS (GTPγS condition).

For experiments using purified proteins as source of soluble binding ligands, the following His-tagged proteins were used (range of protein amounts used in different experiments is in parenthesis): rat His-Gαi3 (0.3-14 μg), human His-Gαi3 (8 μg), His-GINIP (1-12 μg), His-Gαi1 (1 μg), His-Gαi2 (1 μg), His-Gαo (1 μg), His-RGS4 (0.2 μg). Aliquots of protein stored at −80 °C were quickly thawed and cleared by centrifugation at 14,000 × *g* for 2 minutes before addition to tubes containing the GST-fused proteins immobilized on GSH-agarose beads in a final volume of 400 μl. Tubes were incubated for 4 h at 4 °C with constant rotation. Beads were washed three times with 1 ml of wash buffer (4.3 mM Na_2_HPO_4_, 1.4 mM KH_2_PO_4_, pH 7.4, 137 mM NaCl, 2.7 mM KCl, 0.1% (v/v) Tween 20, 10 mM MgCl_2_, 5 mM EDTA, 1 mM DTT) supplemented with GDP, GDPꞏAlF_4_^−^ or GTPγS as indicated above, and resin-bound proteins were eluted with Laemmli sample buffer by incubation at 65 °C for 10 min. Proteins were separated by SDS-PAGE and immunoblotted with antibodies as indicated under “*Protein Electrophoresis and Immunoblotting*.”

For experiments using lysates of cultured cells as a source of soluble binding ligands, HEK293T cells (ATCC CRL-3216) were grown at 37 °C, 5% CO_2_ in DMEM (Gibco, 11965-092) supplemented with 10% fetal bovine serum (Hyclone, SH30072.03), 100 units/ml penicillin, 100 μg/ml streptomycin, and 2 mM L-glutamine (Corning, 30-009-CI). Approximately two million HEK293T cells were seeded on 10-cm dishes and transfected the day after using the calcium phosphate method with plasmids encoding the following constructs (DNA amounts in parentheses): Gαi3-FLAG (6 μg), Gαz-EE (6 μg), Gαo (6 μg), Gαs (6 μg), Gαq-HA (6 μg), Gα12-MYC (6 μg). Cell medium was changed 6 h after transfection, and approximately 24 h later, cells were lysed at 4 °C with 700 μl of lysis buffer (20 mM HEPES, pH 7.2, 125 mM K(CH_3_COO), 0.4% (v/v) Triton X-100, 1 mM DTT, 10 mM β-glycerophosphate, and 0.5 mM Na_3_VO_4_ supplemented with a SigmaFAST protease inhibitor mixture (Sigma, cat# S8830)). Cell lysates were cleared by centrifugation at 14,000 × *g* for 10 minutes, and supplemented with GDP or GDPꞏAlF_4_^−^ as indicated above. One hundred microliters (~400 μg of total protein) of these cell lysates were added to the GST-fused proteins immobilized on GSH-agarose beads in a final volume of 400 μl. Tubes were incubated for 4 h at 4 °C with constant rotation, then washed and eluted according to the protocol described above for purified soluble ligands.

For experiments using brain tissue as a source of endogenous GINIP as binding ligand, the lysates were prepared as follows. Whole brains of adult mice were isolated and lysed in one ml (per brain) of modified RIPA buffer (20 mM Tris-HCl pH 7.4, 150 mM NaCl, 1% (v/v) Nonidet P-40, 1% (w/v) sodium deoxycholate, 0.1% (w/v) SDS, 0.1% (v/v) Triton X100 supplemented with SigmaFAST protease inhibitor mixture) by grinding using tissue homogenizer (Fisherbrand, 15340167) on ice. Lysates were cleared by centrifugation at 14,000 × *g* for 10 minutes at 4 °C twice, aliquoted and stored at −80 °C as single use aliquots for experiments to avoid freeze/thaw cycles. After thawing, lysates were cleared by centrifugation at 14,000 × *g* for 10 minutes at 4 °C. Twenty-five μl of brain lysate (~500 μg of total protein) were added to GST-fused proteins immobilized on GSH-agarose beads in volume of 300 μl of binding buffer supplemented with GDP or GDPꞏAlF_4^−^_ as indicated above. Tubes were incubated for 4 h at 4 °C with constant rotation, then washed according to the protocol described above for purified soluble ligands and eluted with Laemmli sample buffer by incubation at 37 °C for 10 min.

### Protein electrophoresis and immunoblotting

Protein samples were prepared in Laemmli sample buffer as described in previous sections. For the analysis of GINIP expression in mouse tissues, samples were prepared using the same buffer and procedure as for the preparation of brain lysates as described in “*Pulldown assays.*” Proteins were separated by SDS-PAGE and transferred to PVDF membranes, which were blocked with 5% (w/v) nonfat dry milk and sequentially incubated with primary and secondary antibodies. For protein-protein-binding experiments with GST-fused proteins, PVDF membranes were stained with Ponceau S and scanned before blocking. The primary antibodies used were the following (dilution in parenthesis): rabbit Gαi3, SCBT #sc-365422 (1:1000); mouse FLAG, Sigma #F1804 (1:1000); rabbit EE (Glu-Glu), Millipore #AB3788 (1:1000); mouse Gαo, SCBT #sc-13532 (1:1000); rabbit Gαs/olf, SCBT #sc-383 (1:1000); mouse HA, Roche #11583816001 (1:1000); mouse MYC, Cell Signaling #2276 (1:1000); mouse His, Sigma #H1029 (1:2500); mouse α-tubulin, Sigma #DM1A (1:2000); rabbit β-actin, LI-COR #926-42212 (1:1000); rabbit GAIP, serum gifted by M. Farquhar (Garcia-Marcos et al., 2011a; Vries et al., 1998)(1:2000); rabbit pan-Gβ, SCBT #sc-166123 (1:250); goat GINIP, SCBT #sc-247284 (1:1000); mouse Gαq, SCBT #sc-393 (1:1000); mouse Tuj1, SCBT #sc80005 (1:2000); mouse GFAP, Merck #MAB360 (1:2000). The secondary antibodies were (dilution in parenthesis): goat anti-rabbit Alexa Fluor 680, Invitrogen #A21077 (1:10,000); goat anti-mouse Alexa Fluor 680, Invitrogen #A21058 (1:10,000); goat anti-mouse IRDye 800, LI-COR #926-32210 (1:10,000); goat anti-rabbit DyLight 800, Thermo #35571 (1:10,000); donkey anti-goat IRDye 680RD, LI-COR #926-68074. Infrared imaging of immunoblots was performed using an Odyssey CLx Infrared Imaging System (LI-COR). Images were processed using ImageJ software (National Institutes of Health) or Image Studio software (LI-COR), and assembled for presentation using Photoshop and Illustrator software (Adobe).

### GTPγS-binding assays

GTPγS-binding assays were performed as described previously ((Garcia-Marcos et al., 2010; Leyme et al., 2014b). Purified His-Gαi3 (100 nM) was diluted in assay buffer (20 mM Na-HEPES, pH 8, 100 mM NaCl, 1 mM EDTA, 25 mM MgCl_2_, 1 mM DTT, 0.05% (w/v) C_12_E_10_) and preincubated with purified proteins (as indicated in the figures) for 15 min at 30 °C. Reactions were initiated by adding an equal volume of assay buffer containing 1 μM [^35^S]GTPγS (~50 cpm/fmol, Perkin Elmer) at 30 °C. Duplicate aliquots (25 μl) were removed after 15 minutes and binding of radioactive nucleotide was stopped by addition of 3 ml of ice-cold wash buffer (20 mM Tris-HCl, pH 8.0, 100 mM NaCl, 25 mM MgCl_2_). The quenched reactions were rapidly passed through BA-85 nitrocellulose filters (GE Healthcare, 10402506) and washed with 4 ml of cold wash buffer. Filters were dried and subjected to liquid scintillation counting. Background [^35^S]GTPγS detected in the absence of G protein was subtracted from each reaction. Data are expressed as percentage of [^35^S]GTPγS binding relative to a control reaction containing the G protein alone (% of control).

### GTPase assays

Assays to measure GTPase activity under steady-state conditions were performed as described previously (Garcia-Marcos et al., 2010; Leyme et al., 2014b). These experiments were carried out with Gαi1 R178M/A326S (His-Gαi1^RM/AS^), a previously described (Zielinski et al., 2009) double mutant that simultaneously increases the rate of nucleotide exchange and decreases GTP hydrolysis. The rate limiting step of GTPase reactions for Gαi1^RM/AS^ is nucleotide hydrolysis (Zielinski et al., 2009) as opposed to the limiting rate of nucleotide exchange with the wild-type protein (Garcia-Marcos et al., 2010; Ross and Wilkie, 2000), which allows to measure changes in nucleotide hydrolysis such as those caused by GAPs under steady-state conditions. His-Gαi1^RM/AS^ (100 nM) was diluted in assay buffer (20 mm Na-HEPES, pH 8, 100 mm NaCl, 1 mm EDTA, 2 mm MgCl_2_, 1 mm DTT, 0.05% (w/v) C_12_E_10_) with purified proteins (as indicated in the figures) at 4 °C. Reactions were initiated by adding an equal volume of assay buffer containing 1 μm [γ-^32^P]GTP (~50 cpm/fmol, Perkin Elmer) at 30 °C. Duplicate aliquots (25 μl) were removed after 30 minutes, and the reaction was stopped by the addition of 975 μl of ice-cold 5% (w/v) activated charcoal in 20 mM H_3_PO_4_, pH 3. Samples were centrifuged for 10 min at 10,000 × *g*, and 500 μl of the resultant supernatants were subjected to liquid scintillation counting to quantify the amount of [^32^P]P_i_ released. Background [^32^P]P_i_ detected in the absence of G protein was subtracted from each reaction. Data are expressed as percentage of [γ-^32^P]P_i_ released relative to a control reaction containing the G protein alone (% of control).

### cAMP measurements in HEK239T cells by BRET

HEK293T cells (ATCC CRL-3216) were grown at 37 °C, 5% CO_2_ in DMEM medium (Gibco, 11965-092) supplemented with 10% fetal bovine serum (Hyclone, SH30072.03), 100 units/ml penicillin, 100 μg/ml streptomycin, and 2 mM L-glutamine (Corning, 30-009-CI). Approximately 400,000 cells/well were seeded on 6-well plates coated with 0.1% gelatin and transfected ~24 hr later using the calcium phosphate method with plasmids encoding the following constructs (DNA amounts in parentheses): Nluc-EPAC-VV (0.05 μg), GABA_B_R1a (0.2 μg), GABA_B_R2 (0.2 μg), α2_A_-AR (0.2 μg), FLAG-D2R (0.2 μg), Gαi3 WT (0.5 μg), and GINIP-FLAG (2 μg). Total DNA amount per well was equalized by supplementing with empty pcDNA3.1 as needed. Cell medium was changed 6 h after transfection, and approximately 16-24 h after transfection, cells were washed and gently scraped in room temperature PBS, centrifuged (5 min at 550 × *g*), and resuspended in BRET buffer (140 mM NaCl, 5 mM KCl, 1 mM MgCl_2_, 1 mM CaCl_2_, 0.37 mM NaH_2_PO_4_, 24 mM NaHCO_3_, 10 mM HEPES and 0.1% glucose, pH 7.4) at a concentration of ~10^6^ cells/ml. ~25,000 cells/well were added to a white opaque 96-well plate (Opti-Plate, PerkinElmer Life Sciences, 6005290) and mixed with the nanoluciferase substrate Nano-Glo (Promega, N1120, final dilution 1:200) before measuring luminescence. Luminescence signals at 460 ± 80 and 535 ± 35 nm were measured at 28 °C every 2 s in a BMG Labtech POLARStar Omega plate reader and BRET was calculated as the ratio between the emission intensity at 535 nm divided by the emission intensity at 460 nm. Since the Nluc-EPAC-VV construct reports cAMP binding as a decrease in BRET, results were processed as the inverse of the BRET ratio (BRET^−1^) to make it more intuitive. After subtraction of a basal signal measured for 30 s before stimulation with forskolin (ΔBRET^−1^), results were normalized to the maximum response of forskolin detected prior to the addition of GPCR agonists (cAMP (normalized ΔBRET^−1^). For calculation of the agonist-mediated inhibition of cAMP induced by forskolin “inhibition (%FSK),” the average of the last six time points of ΔBRET^−1^ curves were used. Baseline ΔBRET^−1^ values from cells not treated with forskolin were subtracted from values obtained from forskolin and agonist treated or forskolin treated conditions. Inhibition (%FSK) was represented as the ratio of forskolin and agonist treated divided by forskolin treated.

### Measurement of Gαi3-AC5 association in HEK293T cells by BRET

HEK293T cells were cultured, transfected, and harvested as described in “*cAMP measurements in HEK239T cells by BRET*.” Cells were transfected with plasmids encoding the following constructs (DNA amounts in parentheses): GABA_B_R1a (0.2 μg), GABA_B_R2 (0.2 μg), α2_A_-AR (0.2 μg), YFP-hAC5 (0.5 μg), Gαi3-Nluc(a/b) WT or Q204L (0.1 μg), GINIP-FLAG (0.1-2 μg). Total DNA amount per well was equalized by supplementing with empty pcDNA3.1 as needed. Luminescence measurements were carried out as in “*cAMP measurements in HEK239T cells by BRET*,” except that signals were recorded every 0.24 s for kinetic measurements of Gαi3-AC5 association upon modulation of GPCR activation. For endpoint measurements comparing the association of Gαi3 WT or Gαi3 Q204L (QL) under steady-state conditions, signals were recorded as an average of 3 measurements 30 s apart. BRET was calculated as the ratio between the emission intensity at 535 nm divided by the emission intensity at 460 nm for both kinetic and endpoint measurements. Results for the kinetic measurements were presented as increase in BRET after subtraction of the basal signal measured for 30 s before GPCR stimulation (ΔBRET (baseline)), whereas for endpoint measurements they were presented as difference in BRET compared with the BRET signal in cells expressing Gαi3-Nluc WT and YFP-hAC5 in the absence of GINIP (ΔBRET (WT control)).

### Free Gβγ measurements in HEK293T cells by BRET

HEK293T cells were cultured, transfected, and harvested as described in “*cAMP measurements in HEK239T cells by BRET*.” Cells were transfected with plasmids encoding the following constructs (DNA amounts in parentheses): VN-Gγ2 (0.2 μg), VC-Gβ1 (0.2 μg), masGRK3ct-Nluc (0.2 μg), Gαi3 WT (1 μg), Gαo WT (1 μg), GABA_B_R1a (0.2 μg), GABA_B_R2 (0.2 μg), α2_A_-AR (0.2 μg), GAIP (0.5 μg), RGS7 (1 μg), Gβ_5_ (1 μg), R7BP (1 μg), RGS8 (2 μg), RGS12 (0.2 μg), GINIP-FLAG (2 μg). Luminescence measurements and BRET calculations were carried out as in “*cAMP measurements in HEK239T cells by BRET*,” except signals were recorded every 0.24 s. Results were presented as increase in BRET after subtraction of the basal signal measured for 30 s before any stimulation (ΔBRET (baseline)). For the calculation of response amplitudes, the difference between the raw BRET ratio before and 60 s after agonist stimulation was calculated. For the presentation and calculation of G protein deactivation rates, the minimum BRET value reached after the addition of antagonist (plateau signal) was subtracted from the raw BRET ratio of each timepoint (recovery corrected ΔBRET), and each resulting value was scaled to as the percentage of the maximal BRET value right before the addition of antagonist (“% Maximum response”). G protein deactivation rate constants (*k*) were determined by fitting the recovery-corrected ΔBRET values after the addition of antagonist to a one-phase decay equation (*Y*=(*Y*_0_ − *Plateau*) × *e*^(−*kX*)^ + *Plateau*) in Prism 9 (Graphpad), where *Y_0_* is the starting value right before the addition of antagonist (constrained to 100) and *Plateau* is the near-zero minimum estimated by the fit.

### Adenylyl cyclase activity *in vitro*

Regulation of adenylyl cyclase (AC) activity by purified G proteins was determined by measuring the production of cAMP upon reconstitution of the enzyme activity using the purified AC5 C1 and AC2 C2 domains as previously described (Dessauer et al., 1998) with modifications. Before AC activity measurements myr-Gαi1 and His-Gαs were loaded with GTPγS by incubating them at 30 °C with 150 μM GTPγS in buffer (20 mM Tris-HCl, 20 mM NaCl, 1 mM MgCl_2_, 1 mM DTT, 5 % glycerol (v:v)) for 3 hours or 45 minutes, respectively. GTPγS-loaded G proteins were aliquoted and stored at −80 °C. AC reactions were carried out in technical duplicates in a final volume of 40 μl, and all reactants were diluted in assay buffer (50 mM HEPES, 2 mM MgCl_2_, 1 mM EDTA, 0.5 mg/mL BSA, 1 mM DTT, pH 8.0) unless otherwise indicated. 8 μl of 250 nM AC5 C1, 8 μl of 5X concentrated stocks of myr-Gαi1-GTPγS (or the same volume of assay buffer without Gαi1), and 8 μl of 10 μM GST-GINIP (or the same volume of PBS for conditions without GINIP) were mixed and incubated on ice for 15 minutes. Simultaneously, a stock of 1.66 μM AC2 C2 and 0.33 μM Gαs-GTPγS was mixed and incubated on ice for 15 minutes. 12 μl of AC2 C2/ Gαs-GTPγS was added to the tubes containing 24 μl AC5 C1, myr-Gαi1-GTPγS, and GST-GINIP. Samples were then incubated for 15 minutes at room temperature before starting the reactions by addition of 4 μl of ATP and MgCl_2_ solution (10 mM ATP, 50 mM MgCl_2_) and rapidly transferring the tubes to a heat block at 30 °C. The final concentrations of reactants were: AC5 C1, 50 nM; myr-Gαi1-GTPγS, 0.25-2 μM; GST-GINIP, 2 μM; AC2 C2, 0.5 μM; Gαs-GTPγS, 0.1 μM; ATP, 1 mM; MgCl_2_, 5 mM. Reactions were stopped after 10 minutes by rapidly transferring the tubes to a heat block at 95 °C for 5 minutes. Samples were centrifuged at 5,000 × *g* for 2 min, and an aliquot from the supernatant was collected to quantify cAMP using the LANCE cAMP kit (Perkin Elmer, cat#AD0262) according to the manufacturer protocol. Time-resolved fluorescence measurements to quantify cAMP were done on a TECAN Infinite M1000 plate reader in white 384-well ProxiPlates (Perkin Elmer, 6008280). Total cAMP was determined based on a standard curve, and each technical duplicate was measured individually before averaging cAMP values. Specific activity was calculated as nmol cAMP / min / mg AC; mg AC was based on the concentration of AC5 C1, the limiting reactant in the C1/C2 complex. AC activity was background corrected by subtracting the signal obtained for AC5 C1 and AC2 C2 in the absence of Gαs-GTPγS. AC activity was then presented as a percentage of Gαs-GTPγS stimulated AC activity.

### Mouse strains and breeding

All animal experiments were carried out in agreement with the institutional guidelines provided by the Institutional Animal Care and Use Committee at Boston University (protocol number PROTO 201800460), per applicable laws and regulations. Heterozygous mice bearing the reporter and null allele GINIP 1a (C57BL/6N-A^tm1Brd^/a ^Phf24tm1a(EUCOMM)Hmgu^/BcmMmucd, Stock# 037754-UCD) were obtained from the Mutant Mouse Resource & Research Centers (MMRRC), and bred by intercrossing to generate cohorts of littermate animals for experiments and to maintain the line. The conditional GINIP flox allele was generated by crossing GINIP 1a/1a homozygous mice with Rosa26 Flpe homozygous mice (B6N.129S4-Gt(ROSA)26Sor^tm1(FLP1)Dym^/J, JAX stock# 016226). The resulting GINIP +/flox; Rosa26 +/Flpe animals were intercrossed to obtain animals bearing the GINIP flox allele but without the Rosa26 Flpe allele. GINIP flox mice were crossed with an Emx1 Cre driver line (B6.129S2-*Emx1*^tm1(cre)Krj^/J, JAX stock# 005628) or a VGAT Cre driver line (B6J.129S6(FVB)-*Slc32a1*^tm2(cre)Lowl^/MwarJ, JAX stock# 028862) to achieve specific GINIP knock-out (GINIP Δ null allele) in excitatory neurons of the telencephalon (Gorski et al., 2002) or in inhibitory neurons (Vong et al., 2011), respectively. To obtain cohorts of littermates for experiments assessing the conditional ablation of GINIP in specific neuron populations, animals with the genotype GINIP flox/flox; Cre/+ were generated first and then crossed with GINIP flox/flox mice. Animal genotyping was carried out by PCR of genomic DNA extracted from tail clipping using a three-primer system. To distinguish between wild-type and 1a alleles, a forward primer targeting the intronic region between exons 5 and 6 (Fw.Intron5: TTAAAGTTGCACAACCCACTAGAAGC) and a forward primer targeting the gene trap cassette (Fw.1aTrap: GGGATCTCATGCTGGAGTTCTTCG) were used with a common reverse primer targeting the intronic region between exons 5 and 6 (Rv.Intron5: GAGCTGAGTGACTCTAGGGATGAACC) resulting in a 308 bp band for the wild-type allele and a 584 bp band for the 1a allele. The Fw.Intron5 and Rv.Intron5 primer set also allows for detection of the GINIP flox allele as 513 bp amplicon. Emx1 Cre (Yoo et al., 2019) and VGAT Cre (Weaver et al., 2018) lines were genotyped as previously described. To detect GINIP Δ null allele obtained after Cre-mediated recombination of the GINIP flox allele, the Fw.Intron5 primer described above was used with a reverse primer targeting downstream of exon 10 (Rv.5UTR: CCAGGCAGAAATCCACAAACTAGG) resulting in a 531 bp band.

### β-galactosidase staining

GINIP +/1a and GINIP +/+ littermates (3 month-old) were anesthetized and transcardially perfused with cold PBS. Brains were rapidly removed from the skull, and placed in an acrylic matrix (Electron Microscopy Sciences, 69080-C or 69080-S) to make either coronal or sagittal slices (1 mm thickness). Slices were rinsed twice with PBS and incubated in staining solution (0.1 M phosphate buffer, pH 7.3, 20 mM Tris HCl, 2 mM MgCl_2_, 0.01% sodium deoxycholate, 0.02% NP-40, 5 mM potassium ferricyanide, 5 mM potassium ferrocyanide, 1 mg/ml X-gal (5-bromo-4-chloro-3-indolyl-β-D-galactopyranoside)) at room temperature with gentle rocking for 1 hour. Staining was stopped by removal of the staining solution, and samples were washed with PBS 3 times, followed by fixation in 4% (w/v) paraformaldehyde in PBS at room temperature for 30 min. Slices were moved to the stage of an Olympus SZX16 stereo microscope equipped with a digital camera (U-TV0.63XC) and with a SDF PLAPO 0.5XPF objective lens and imaged by bright-field microscopy at room temperature using QImaging (Teledyne) software. Individual images were assembled for presentation in Photoshop and Illustrator software (Adobe).

### Fluorescence mRNA *in situ* hybridization (RNAScope®)

Mice (4-5 month-old, genotype-matched littermates) were anesthetized and transcardially perfused with cold PBS. Brains were rapidly removed from the skull, submerged in cryo-embedding medium (OCT) (Fisherbrand, 23730571), and frozen in dry-ice prior to storage of the OCT tissue blocks at −80 °C. Blocks were equilibrated to −20 °C and sectioned in a HM 550 VP cryostat microtome (MICROM GmbH, 956444). Twenty μm sections were mounted onto Superfrost Plus slides (Fisherbrand, 22037246) and stored at −80 °C in air-tight, light-proof containers until use. Frozen slides were placed in a Tissue Tek Slide Rack (StatLab, LWS2124), and immediately immersed in pre-chilled fixative solution (10% neutral buffered formalin, Fisherbrand, 305510) for 15 min at 4 °C. After fixation, sections were washed at room temperature with PBS twice by dipping the rack in the solution container 3-5 times during a period of ~3 min for each wash. Sections were dehydrated by sequential 5-minute incubations at room temperature in 50%, 75% and 100% (v/v) ethanol. Slides were air-dried by incubation at room temperature for approximately 5 min before drawing a hydrophobic barrier around each section with an Immedge™ hydrophobic barrier pen (Vector, H-4000). Subsequent steps were carried out using the RNAscope 2.0 Assay kit following the manufacturer’s instructions (Bio-techne, 320850). Briefly, ~5 drops of Protease IV solution (Bio-techne, 322340) were added to each section, followed by incubation at room temperature for 30 min. Slides were washed with PBS twice by dipping the slide rack in the solution container 3-5 times during a period of ~3 min for each wash. After removing the excess of liquid by flicking, the Amp 1-FL reagent containing the hybridization probe(s) for GINIP (Mm-Phf24-O1), vGlut1 (Mm-Slc17a7-C2) and/or VGAT (Mm-Slc32a1-C3), all from Bio-techne, was added to cover the slides, followed by a 30 min incubation at 40 °C in a HybEZ™ II hybridization oven (Bio-techne, 240200ACD). After two washes with 1X Wash buffer (Bio-techne, 310091) slides were sequentially incubated in 100 μl of the following reagent solutions containing the signal amplifiers (Amp 2-FL and Amp 3-FL) and the fluorescent tracers (Amp 4-FL) for each mRNA probe at 40 °C in a HybEZ™ II hybridization oven for 30 min, 15 min, and 30 min, respectively. Samples were washed twice with 1X Wash buffer after each incubation. The slides were stained with DAPI for 1 minute (Bio-techne, 320858), and mounted in ProLong Diamond Antifade (Invitrogen, P36965). Mounted slides were cured overnight at room temperature prior to imaging. Stained sections were imaged by wide-field microscopy at room temperature using a Zeiss Axio Observer Z1 microscope equipped with a digital camera (C10600/ORCA-R2 Hamamatsu Photonics). Images were taken with a 63x oil-immersion objective (NA = 1.4; working distance = 0.19 mm) using ZEN software. Individual images were assembled for presentation in Photoshop and Illustrator software (Adobe). The custom-made probe for GINIP mRNA (Mm-Phf24-O1, Bio-techne) recognizes the region from nucleotide 857 to 1491 of the mouse GINIP mRNA sequence NM_001346526.1. This region is present in mRNA from the GINIP wild-type and GINIP flox alleles, but not in the GINIP 1a allele or the GINIP flox allele after Cre-mediated recombination (GINIP Δ null allele).

### Mouse primary cortical cultures

Cortical neuron cultures were established from neonatal mouse brains (wild-type C57BL/6, Charles River, strain code 027, or the genotype(s) indicated for each experiment) as previously described (Kaech and Banker, 2006) with modifications. Newborn mouse pups (P0) were euthanized by decapitation, and brains rapidly placed in cold HBSS (Corning, 21-022-CV) after removal from the skull. The cerebrum was detached from other brain regions under a stereomicroscope by removal of the olfactory bulb and cerebellum, and the cortex dissected out with forceps. Tissue was minced into approximately 1-2 mm pieces using a sterile razor blade, and digested with 0.05% Trypsin in HBSS for 10 min at 37°C. Trypsinized tissue was washed three times with HBSS by cycles of gravity sedimentation of tissue and aspiration of buffer. Washed tissue was resuspended in DMEM supplemented with 10% FBS, 100 U/ml penicillin, 100 μg/ml streptomycin, and 2 mM L-glutamine (complete DMEM) before passing through a sterile 40 μm cell strainer (Fisherbrand, 22363547). Cells were counted and seeded on poly-L-lysine coated plates or coverslips. Coating was performed overnight at room temperature with 0.1 mg/mL poly-L-lysine hydrobromide (Sigma, P1955), followed by 3 washes with HBSS and addition of complete DMEM before seeding. Four hours after seeding, half of the medium was replaced by Neurobasal media (GIBCO, 21103049) with B-27 supplement (GIBCO, 17504001) and 1x Glutamax-I (GIBCO, 35050061) (complete neural medium). On day in vitro 3 (DIV3), one half of the media was replaced with complete neural media supplemented with 5 μM AraC to block glial cell proliferation. Beginning DIV5, half of the media was replaced by fresh complete neural medium every other day. For experiments comparing the expression of GINIP in neuron cultures and glial cell cultures by immunoblotting, 500,000 cells were seeded on 60 mm dishes and cultured as described above, except that for the glial cultures AraC was not added at DIV3. At DIV12, neuron or glial cells were washed twice by cold PBS and harvested by scraping followed by centrifugation at 10,000 x *g* at 4 °C. Cell pellets were lysed by resuspending with lysis buffer(20 mM HEPES, pH 7.2, 125 mM K(CH_3_COO), 0.4% (v/v) Triton X-100, 1 mM DTT, 10 mM β-glycerophosphate, and 0.5 mM Na_3_VO_4_ supplemented with a SigmaFAST protease inhibitor mixture) and cleared by centrifugation at 14,000 × *g* for 10 minutes at 4°C. Proteins concentration was quantified by Bradford and lysates boiled for 5 min in Laemmli sample buffer before “*Protein Electrophoresis and Immunoblotting*.”

### Immunofluorescence

Cortical neuron cultures were established from neonatal mouse brains as described in “*Mouse primary cortical cultures.*” 50,000 cells were plated on poly-L-lysine coated 12 mm coverslips in 24 well plates. DIV12 cells were used for experiments evaluating neuronal versus glial marker expression, whereas DIV21 cells were used for experiments evaluating the presence of different synaptic markers in dendrites. Cells were washed with PBS twice for 2 minutes at room temperature and fixed with 4% (w/v) PFA in PBS for 15 minutes at room temperature. After 2 additional washes with PBS, cells were permeabilized and blocked at room temperature for 1 hour in blocking buffer (5% (w/v) Bovine Serum Albumin (BSA), 5% (v/v) goat serum, 0.1% (v/v) Triton X-100 in PBS). Coverslips were placed upside down over 25 µl of buffer (1% (w/v) BSA, 1% (v/v) goat serum, 0.1% (v/v) Triton X-100 in PBS) spotted on parafilm containing primary antibodies in the following dilutions: rabbit GINIP, Aviva ARP70657_P050 (1:100); mouse Tuj1, SCBT sc-80005 (1:200); mouse GFAP Merck MAB360 (1:200); mouse synaptophysin, SCBT sc-55507 (1:200); mouse vGlut1, SCBT sc-377425 (1:50); mouse GAD65, DSHB AB_528264 (1:50); mouse PSD95, ABcam ab13552 (1:200); mouse Gephyrin, Synaptic Systems 147011 (1:200). After overnight incubation at 4°C in a humid chamber, coverslips were washed with PBS 3 times for 5 minutes at room temperature, placed upside down over 30 µl of buffer (1% (w/v) BSA, 1% (v/v) goat serum, 0.1% (v/v) Triton X-100 in PBS) spotted on parafilm containing secondary antibodies in the following dilutions: goat anti-mouse Alexa Fluor 488, Life Technologies A11017 (1:400); goat anti-rabbit Alexa Fluor 488, Life Technologies A11070 (1:400); goat anti-mouse Alexa Fluor 594, Life Technologies A11020 (1:400), and goat anti-rabbit Alexa Fluor 594, Life Technologies A11072 (1:400). After incubation for 1 hour at room temperature in a humid chamber, coverslips were washed with PBS 3 times for 10 minutes at room temperature, followed by staining with DAPI (1:10,000) for 5 minutes at room temperature. After one wash with PBS, coverslips were mounted with ProLong Diamond Antifade (Invitrogen, P36970) and cured overnight at room temperature prior to imaging.

Wide-field microscopy imaging was performed at room temperature with a Zeiss Axio Observer Z1 microscope equipped with a digital camera (C10600/ORCA-R2 Hamamatsu Photonics). Images were taken with a 63x oil-immersion objective (NA = 1.4; working distance = 0.19 mm) using ZEN software. Confocal microscopy imaging was carried out at room temperature with a Zeiss LSM 700. Single sections of confocal images of 0.321 μm thickness along the z axis were taken with a 63x oil-immersion objective (NA 1.4, working distance 0.19 mm) using ZEN software.

For experiments to determine the expression of GINIP in neuron versus glia (**Fig. 5**), the number of NeuN (neuronal marker) or GFAP (astrocyte marker) positive cells that were also positive for GINIP was quantified as a percentage from 6 randomly chosen fields of three independent cultures at DIV12 (2 fields/ experiment). For experiments to determine the subcellular localization of GINIP (**Fig. 6**), the number of GINIP positive puncta (0.25-0.75 µm^2^ area) on dendrites that were also positive for each synaptic marker was quantified as a percentage from 3-4 fields per experiment. Three independent experiments corresponding to separate cultures at DIV21 were quantified. Individual images were assembled for presentation in Photoshop and Illustrator software (Adobe).

### Chemically induced seizures

Animals (n = 11 – 14 per group; approximately 5 – 6 months old; males and females) were assessed prior to start of experiment for any health issues and were individually habituated to sterile empty housing cages prior to start of study. During the study, all subjects were injected with bicuculline (i.p. Bio-Techne Co., Minneapolis, MN USA) at increasing doses (0, 1, 2 and 3 mg/kg), immediately placed into sterile empty housing cage, and tracked (Logitech C920 HD Pro; Logitech International S.A., Newark, CA USA. Canon VIXIA HF MF80 HD; Canon USA, Melville, NY USA) for 30 minutes. Behavior was scored by intensity of 0 to 4 (0 = normal behavior, 1 = wild running, 2 = tonic seizure, 3 = clonic seizure, 4 = cardiac arrest/death) and by number of seizure episodes. The genotypes of all animals were blinded prior to start of study.

### Lentivirus packaging and transduction of primary cortical neurons

Lentiviruses used for the transduction of cultured neurons were concentrated after large scale packaging as described next. HEK293T cells (Lenti-X 293T, Cat# 632180, Takara Bio) were plated on 150 mm diameter dishes (~2.5 million cells / dish) and cultured at 37°C, 5% CO2 in DMEM supplemented with 10% FBS, 100 U/ml penicillin, 100 μg/ml streptomycin, and 2 mM L-glutamine. After 16-24 h, cells were co-transfected with plasmids encoding Gβγ-BERKY1 (pLenti-hSyn-Gβγ-BERKY1) (27 μg / dish), or Nluc-EPAC-VV (pLenti-hSyn-Nluc-EPAC-VV) (27 μg / dish) along with the packaging plasmid psPAX2 (18 μg / dish) and the envelope plasmid pMD2.G (11.25 μg / dish) using the polyethylenimine (PEI) method (Longo et al., 2013) at a 2:1 PEI:DNA ratio. Approximately 16 h after transfection, media was replaced with serum-free media. Lentivirus containing medium was collected 24 h and 48 h after the initial media change (a total of ~70 ml per dish and 4 dishes for each construct). Media was centrifuged for 5 min at 900 x *g* and filtered through a 0.45 μm sterile PES filter (Fisherbrand, cat# FB12566505). Filtered media was centrifuged for overnight (~16 h) at 17,200 x *g* at 4 °C (Sorvall RC6+, ThermoScientific F12-6x500 LEX rotor) to sediment lentiviral particles. Pellets were washed and gently resuspended in 1 ml of PBS and centrifuged at 50,000 × *g* for 1 h at 4°C (Beckman Optima MAX-E, TLA-55 rotor). The resulting pellets were resuspended in 500 μl of PBS to obtain concentrated lentiviral stocks that were stored at −80°C in aliquots. Each aliquot was thawed only once for subsequent experiments.

### Measurement of free Gβγ in primary cortical neurons by BRET

Cortical neuron cultures were established from neonatal brains of GINIP flox/flox mice as described in “*Mouse primary cortical cultures.*” For experiments aimed at measuring the release of free Gβγ in neurons, 100,000 cells were seeded on poly-L-lysine coated 5 mm coverslips in 96 well plates. On DIV6, primary cortical neurons were transduced with AAV encoding Cre recombinase (AAV-hSyn-Cre, Addgene 105553-AAV1) by replacing half of the volume of media in the well with media containing the AAV (final virus dilution 1:500 in 200 μl per well). Two hours later, 100 μl of media were replaced by fresh complete neural medium. Controls were subjected to the same media changes but omitting the AVV. At DIV8, all conditions were transduced with lentiviruses encoding Gβγ-BERKY1 by replacing one half of the media with complete neural media containing lentivirus (final lentivirus dilution 1:200 in 200 μl per well). Two hours after addition of the lentivirus, one half of the medium was replaced with fresh complete neural media and culturing carried on as described in “*Mouse primary cortical cultures*”. BRET measurements were carried out between DIV12 and DIV16. For this, coverslips were transferred to white opaque 96-well plate (Opti-Plate, PerkinElmer Life Sciences) containing 100 μl of BRET buffer (140 mM NaCl, 5 mM KCl, 1 mM MgCl_2_, 1 mM CaCl_2_, 0.37 mM NaH_2_PO_4_, 24 mM NaHCO_3_, 10 mM HEPES and 0.1% glucose, pH 7.4) with nanoluciferase substrate Nano-Glo (Promega, final dilution 1:200). Luminescence signals at 460 ± 80 and 535 ± 35 nm were measured at 28 °C every 0.96 s in a BMG Labtech POLARStar Omega plate reader, and BRET was calculated as the ratio between the emission intensity at 535 nm divided by the emission intensity at 460 nm. Results were calculated as increase in BRET after subtraction of the basal signal measured for 30 s before any stimulation, followed by normalization to the maximum increase in BRET observed in control neurons (from GINIP flox/flox mice without Cre recombination) observed upon stimulation with saturating concentration of the agonist for the GPCR under investigation (insert here how this is called in the Y axis of the corresponding result).

### Measurement of cAMP in primary cortical neurons by BRET

Cortical neuron cultures were established from neonatal brains of GINIP flox/flox mice as described in “*Mouse primary cortical cultures.*” For experiments aimed at measuring intracellular cAMP levels cells were processed as described above in “*Measurement of free Gβγ in primary cortical neurons by BRET*”, except that the lentiviral transductions were carried out with particles for the expression of the BRET-based cAMP biosensor Nluc-EPAC-VV. BRET measurements were carried out as described in *“Measurement of free Gβγ in primary cortical neurons by BRET.”* Luminescence signals at 460 ± 80 and 535 ± 35 nm were measured at 28 °C every 5 seconds in a BMG Labtech POLARStar Omega plate reader and BRET was calculated as the ratio between the emission intensity at 535 nm divided by the emission intensity at 460 nm. Since the Nluc-EPAC-VV construct reports cAMP binding as a decrease in BRET, results were processed as the inverse of the BRET ratio (BRET^−1^) to make it more intuitive. Results are presented as the change in BRET (ΔBRET^−1^) relative to the BRET baseline measured for 30 s before stimulation with forskolin.

### Brain Slice Electrophysiology

Brains from adult (2-5 month-old) GINIP +/+ and GINIP 1a/1a littermate mice were sectioned fresh to generate coronal slices (220 μm thickness) using a Leica VT1000S vibratome (Leica Microsystems). Mice were anesthetized, perfused and decapitated in accordance with national and institutional guidelines. Perfusion was carried out with modified artificial cerebral spinal fluid (mACSF) containing (in mM): 92 NMDG, 20 HEPES, 25 glucose, 30 NaHCO3, 1.2 NaH_2_PO_4_, 2.5 KCl, 5 L-ascorbic acid, 3 sodium pyruvate, 2 thiourea, 10 MgSO_4_, 0.5 CaCl_2_, 12 N-acetyl-l-cysteine, 300−310 mOsm, pH 7.3−7.4 adjusted with 4N HCl. Brain slices were sectioned in cold mACSF, recovered in the same buffer at 32 °C for 10 min, and further transferred to a holding ACSF solution containing (in mM): 92 NaCl, 20 HEPES, 25 glucose, 30 NaHCO_3_, 1.2 NaH_2_PO_4_, 2.5 KCl, 5 L-ascorbic acid, 3 sodium pyruvate, 2 thiourea, 1 MgSO_4_, 2 CaCl_2_, 12 N-acetyl-l-cysteine, 300−310 mOsm, at pH 7.3−7.4 at room temperature to recover for 1h before electrophysiological recordings. All solutions were continuously saturated with 95% O_2_ and 5% CO_2_. Slices were continuously perfused (2 ml/min) during electrophysiological recordings with carbogen saturated ACSF containing (in mM): 125 NaCl, 2.5 KCl, 1.25 NaH_2_PO_4_, 1 MgCl_2_, 26 NaHCO_3_, 11 glucose, 2.4 CaCl_2_, 300− 310 mOsm, at pH 7.3−7.4 at 32°C. Pyramidal neurons from the somatosensory cortex (layer III/IV) were visualized with infrared differential interference contrast video microscopy (FV3000, Olympus) under 40x water-immersion lens and recorded in the whole-cell configuration using a MultiClamp 700B amplifier (2 kHz low-pass Bessel filter and 10 kHz digitization) with pClamp 11 software (Molecular Devices). Patch pipettes were pulled from OD/ID 1.5/0.86 mm capillary glass in a horizontal pipette puller (Model P-1000, Sutter Instrument). Series resistance and whole cell capacitance were monitored during the experiments and recordings with access resistance changes >50% were discarded.

Potassium currents were recorded in the presence of 1 μM tetrodotoxin (TTX) in the bath solution and with the borosilicate glass electrodes (3−5 MΩ) filled with an internal solution containing (in mM): 120 mM potassium gluconate, 20 KCl, 0.05 EGTA, 10 HEPES, 1.5 MgCl_2_, 2.18 Na_2_-ATP, 0.38 Na-GTP, 10.19 sodium phosphocreatine, 275−285 mOsm, and pH 7.3−7.4. Holding current was recorded in voltage clamp mode at a corresponding resting membrane potential of −55-65 mV. Started 10 min after stable whole cell access was obtained, K+ currents were induced by 50 μM baclofen followed by wash with ACSF. Change (Δ) in holding current (pA) after baclofen addition was presented as mean ± S.E.M. from 14 cells per group (representing at least 4-5 animals per group).

Miniature inhibitory postsynaptic currents (mIPSCs) were recorded by adding 1 μM tetrodotoxin (TTX), 10 μM 6-cyano-7-nitroquinoxaline-2,3-dione (CNQX), and 10 μM d-2-amino-5 phosphonopentanoic acid (d-AP5) in the bath solution, and inhibiting K^+^ channels by filling the electrodes with internal solution containing (in mM): 140 mM cesium gluconate, 5 NaCl, 2 MgCl_2_, 10 HEPES, 0.1 EGTA, 2 Na_2_-ATP, 0.4 Na-GTP, 275−285 mOsm, at pH 7.3−7.4. The holding potential was set to +10 mV and current was recorded in voltage clamp mode. Ten min of stable baseline was confirmed after obtaining whole cell access. Different concentrations of baclofen (0.1, 1 and 10 μM) were perfused for 5 min, followed by wash with ACSF. Positive-going outward IPSCs were separately detected at the same baseline using Clampfit analysis software and a template was constructed by averaging >50 mIPSCs for automatic event frequency detection with amplitude above threshold of 5 pA. At least a total of 14 cells from 4-5 different animals were recorded per group. Data were reported as mean ± SEM.

