## Supplemental information for "Fine-tuning GPCR-mediated neuromodulation by biasing signaling through different G-protein subunits"

Contents: Six supplementary figures and their legends
Figure S1-S6

### FIGURE S1

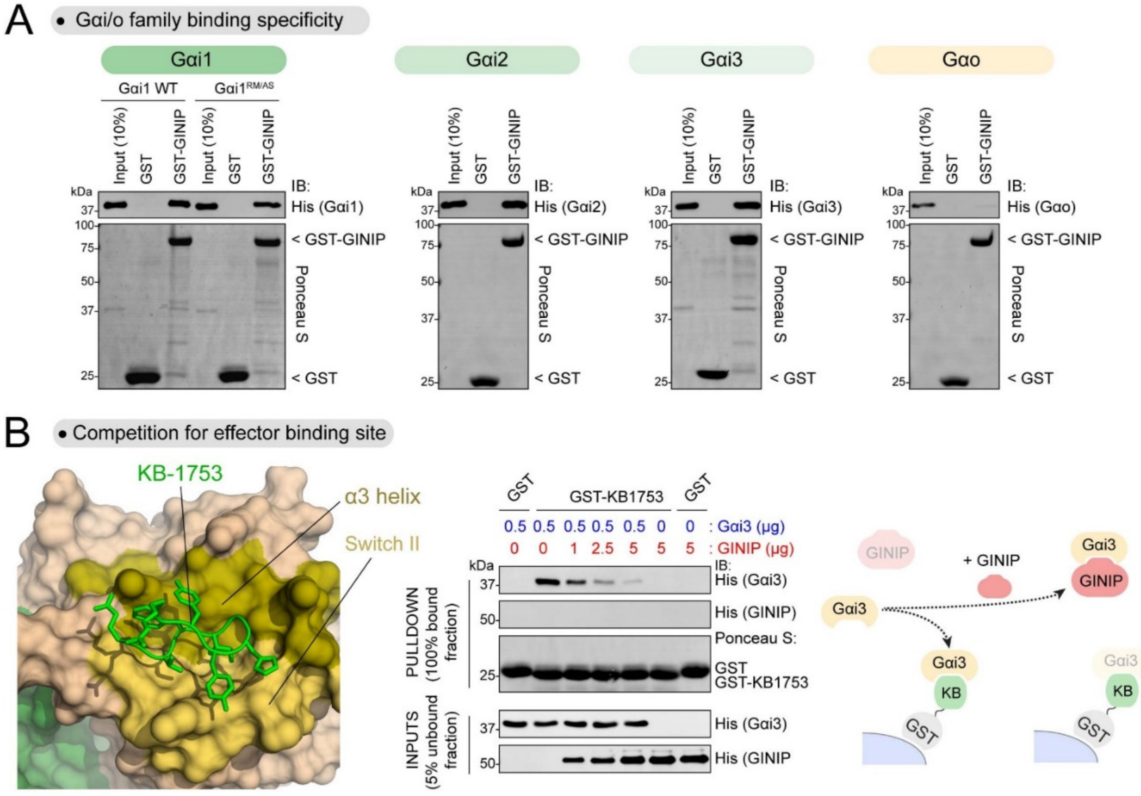


**Figure S1.** **GINIP binds to all Gαi isoforms and competes with the effector-like KB-1753 peptide for binding to Gαi.**

**(A)** GINIP binds similarly to Gαi1/2/3 and Gαi1 R178M/A326S (Gαi1$^{RM/AS}$), but not to Gαo. Purified His-tagged Gαi1 WT, Gαi1$^{RM/AS}$, Gαi2, Gαi3, and Gαo were incubated with GST or GST-GINIP immobilized on glutathione-agarose beads in the presence of GDP·$AlF_4^-$. Bead-bound proteins were detected by Ponceau S staining or by immunoblotting (IB).

**(B)** GINIP competes with the KB-1753 peptide for binding to Gαi. *Left,* Structural model of the Gαi1/KB-1753 complex (PDB: 2G83). *Right,* increasing amounts of purified His-GINIP and a fixed amount of His-Gαi3 loaded with GDP·$AlF_4^-$ were incubated with GST or GST-KB-1753 immobilized on glutathione-agarose beads. Bead-bound (“Pulldown”) and unbound (“Inputs”) fractions were detected by Ponceau S staining or by immunoblotting (IB).

Results are representative of n ≥ 3 experiments.

FIGURE S2

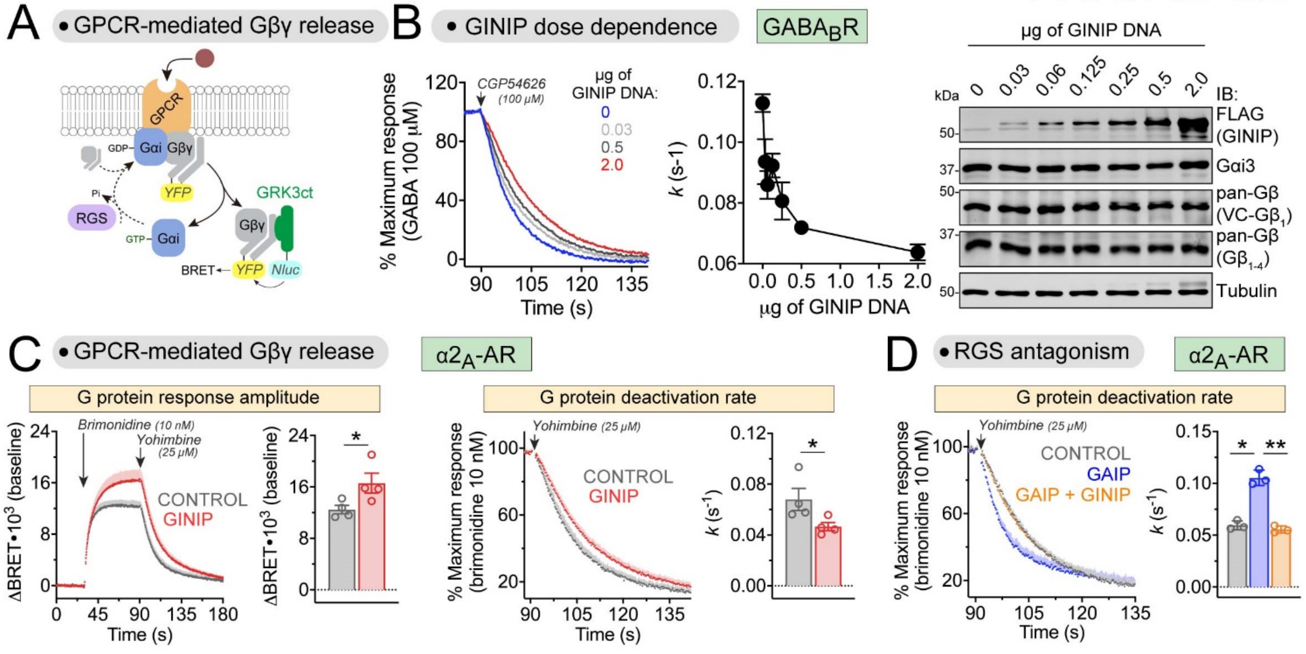


**Figure S2. GINIP modulates Gβγ signaling triggered by different GPCRs.**

**(A)** Diagram of G protein activation/deactivation cycle and BRET-based detection of free Gβγ.

**(B)** DNA dose-dependent effect of GINIP on Gβγ deactivation kinetics triggered by GABA$_B$R. BRET was measured in HEK293T cells expressing the GABA$_B$R in the absence (blue) or presence (gray/red) of increasing amounts of GINIP plasmid corresponding to the kinetic traces. *Left,* Cells were treated with CGP54626 as indicated one minute after treatment with GABA (1 µM). *Center,* G protein deactivation rates were determined by normalizing the BRET data to maximum response and fitting the post-antagonist data to an exponential decay curve to extract rate constant values (*k*). *Right,* Expression of GINIP was validated by immunoblotting (IB). Mean ± S.E.M., n=5.

**(C)** GINIP enhances Gβγ-mediated signaling amplitude and deactivation kinetics triggered by α2-AR. *Left,* BRET was measured in HEK293T cells expressing the α2-AR in the absence (black) or presence (red) of GINIP. Kinetic traces correspond to cells expressing no GINIP (black) or transfected with 2 µg of GINIP plasmid (red). Cells were treated with brimonidine and yohimbine as indicated and the amplitude of the BRET responses quantified 1 min after agonist stimulation. *Right,* G protein deactivation rates were determined by normalizing the BRET data to maximum response and fitting the post-antagonist data to an exponential decay curve to extract rate constant values (*k*). Mean ± S.E.M., n=4. *p<0.05, paired t-test.

**(D)** GINIP antagonizes Gβγ regulation by GAIP upon activation of α2-AR. BRET experiments were carried out and analyzed as in (C) in the absence (grey) or presence of GAIP (blue) or GAIP plus GINIP (orange). Mean ± S.E.M., n=3. *p<0.05, **p<0.01, one-way ANOVA corrected for multiple comparisons (Tukey).

FIGURE S3

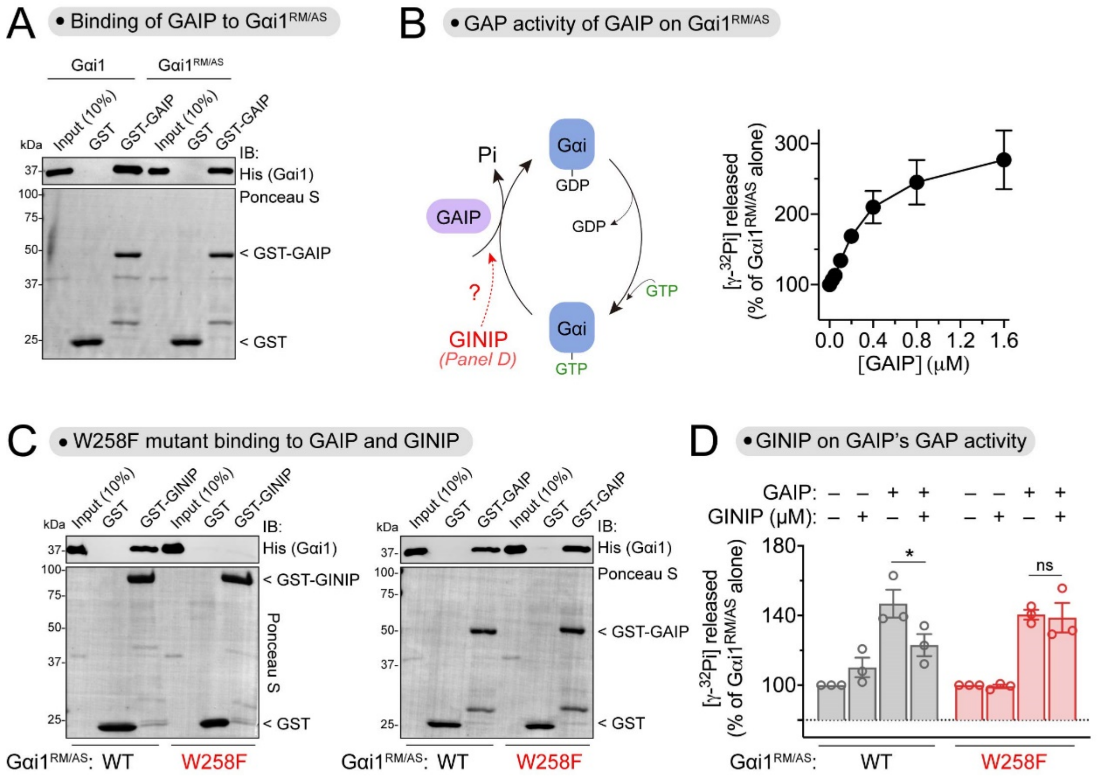


**Figure S3. GINIP antagonizes the GAP activity of GAIP on Gαi *in vitro*.**

**(A)** Gαi1 WT and Gαi1$^{RM/AS}$ bind similarly to GAIP. Purified His-tagged Gαi1 WT or Gαi1$^{RM/AS}$ was incubated with GST or GST-GAIP immobilized on glutathione-agarose beads in the presence of GDP·AlF$_4^-$. Bead-bound proteins were detected by Ponceau S staining or by immunoblotting (IB).

**(B)** Dose-dependent effect of GAIP on steady-state GTP hydrolysis by Gαi1$^{RM/AS}$. Nucleotide hydrolysis was determined by the release of free phosphate (Pi) from GTP in the presence of increasing concentrations of GAIP. Mean ± S.E.M., n=4.

**(C)** W258F mutation disrupts Gαi1$^{RM/AS}$ binding to GINIP but not to GAIP. Purified His-tagged Gαi1$^{RM/AS}$ WT or Gαi1$^{RM/AS}$ W258F was incubated with GST-GINIP (left) or GST-GAIP (right) immobilized on glutathione-agarose beads in the presence of GDP·AlF$_4^-$. Bead-bound proteins were detected by Ponceau S staining or by immunoblotting (IB).

**(D)** GINIP antagonizes the GAP activity of GAIP on Gαi *in vitro*. Nucleotide hydrolysis by Gαi1$^{RM/AS}$ (WT or W258F) was determined in the presence of GAIP and/or GINIP as indicated. Mean ± S.E.M., n=3. ns = not significant, *$p<0.05$, one-way ANOVA corrected for multiple comparisons (Tukey).

FIGURE S4

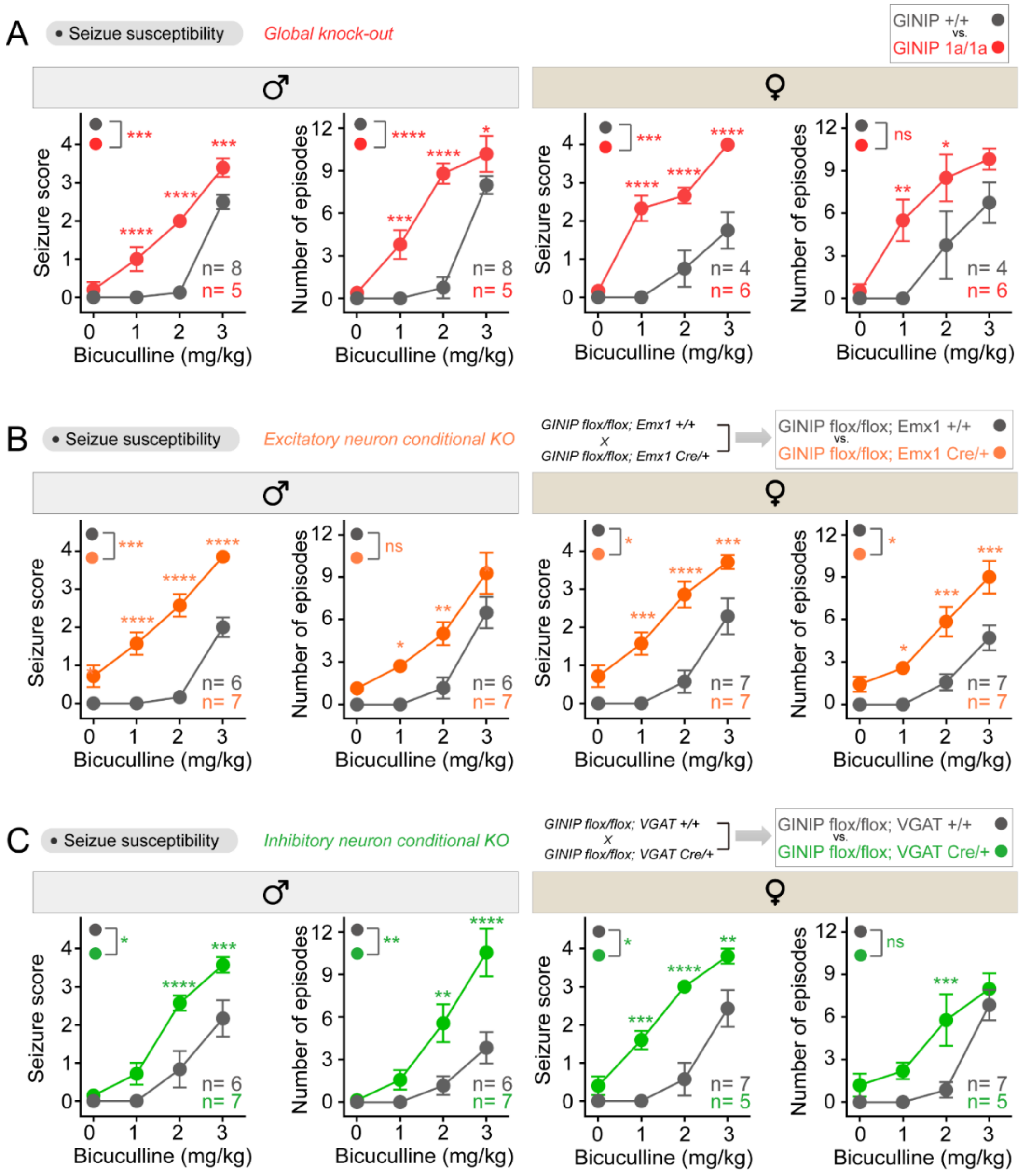


**Figure S4. Loss of GINIP from excitatory and/or inhibitory neurons increases seizure susceptibility in either male or female mice.**

**(A-C)** Seizure susceptibility analysis by sex of global knock-out (KO) (A), GINIP flox/flox; Emx1 Cre/+ (B) and GINIP flox/flox; VGAT Cre/+ (C) mice. GINIP 1a/1a mice were compared to wild-type littermates in (A), whereas GINIP flox/flox mice were compared to littermates bearing an Emx1 Cre driver allele in (B), or a VGAT Cre driver allele in (C). 10-14 week-old mice of each sex were assessed for bicuculline-induced seizure susceptibility by measuring seizure score and number of episodes after bicuculline injection with indicated mice number.

All graphs are mean ± S.E.M. *p<0.05, **p<0.01, ***p<0.001, ****p<0.0001, two-way ANOVA for genotype x concentration of bicuculline, with multiple comparisons at each concentration using Fisher's LSD test.

FIGURE S5

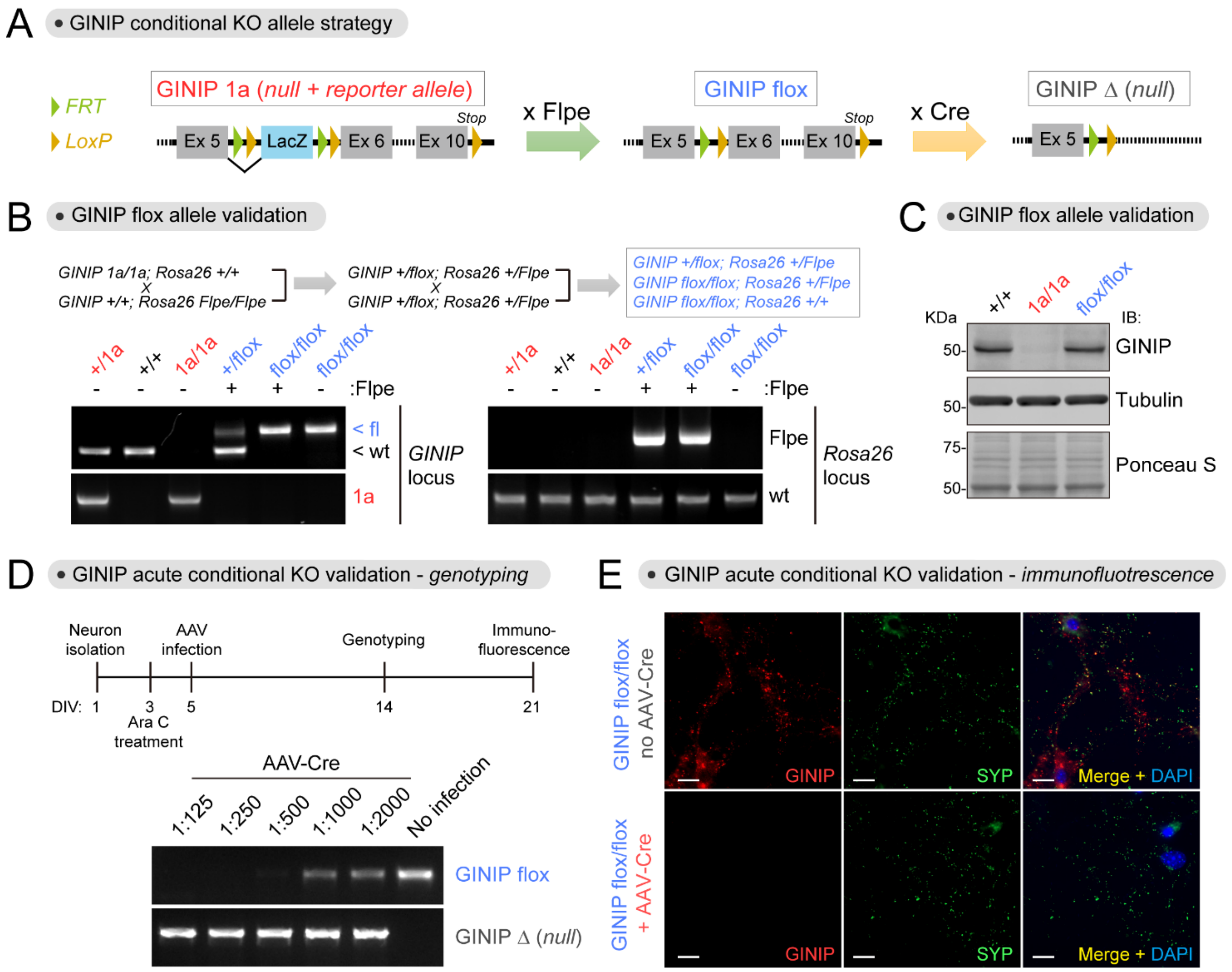


**Figure S5. Establishment and validation of a conditional GINIP null allele (GINIP flox).**

**(A)** Diagram depicting mouse crosses to generate a Cre-dependent conditional GINIP null allele (GINIP flox).

**(B)** Genotyping results of different steps of the crosses carried out to generate the GINIP flox allele.

**(C)** Brains from GINIP flox/flox mice express as much GINIP as in GINIP +/+ mice. Proteins extracted from brains of mice with the indicated genotypes were analyzed by Ponceau S staining or immunoblotting (IB) and indicated. n=3.

**(D, E)** Acute infection of cortical neurons from GINIP flox/flox mice with an AAV-Cre virus leads to ablates GINIP expression. Cortical neurons isolated from GINIP flox/flox mice were infected by AAV-Cre at DIV5 and harvested at DIV14 for genotyping (D) or at DIV21 for immunofluorescence (E). For D, PCR are genotyping results of neurons treated with the indicated dilutions of AAV-Cre virus. For E, neurons were co-stained for GINIP and synaptophysin (SYP). Scale bar = 20 µm. n=3

#### FIGURE S6

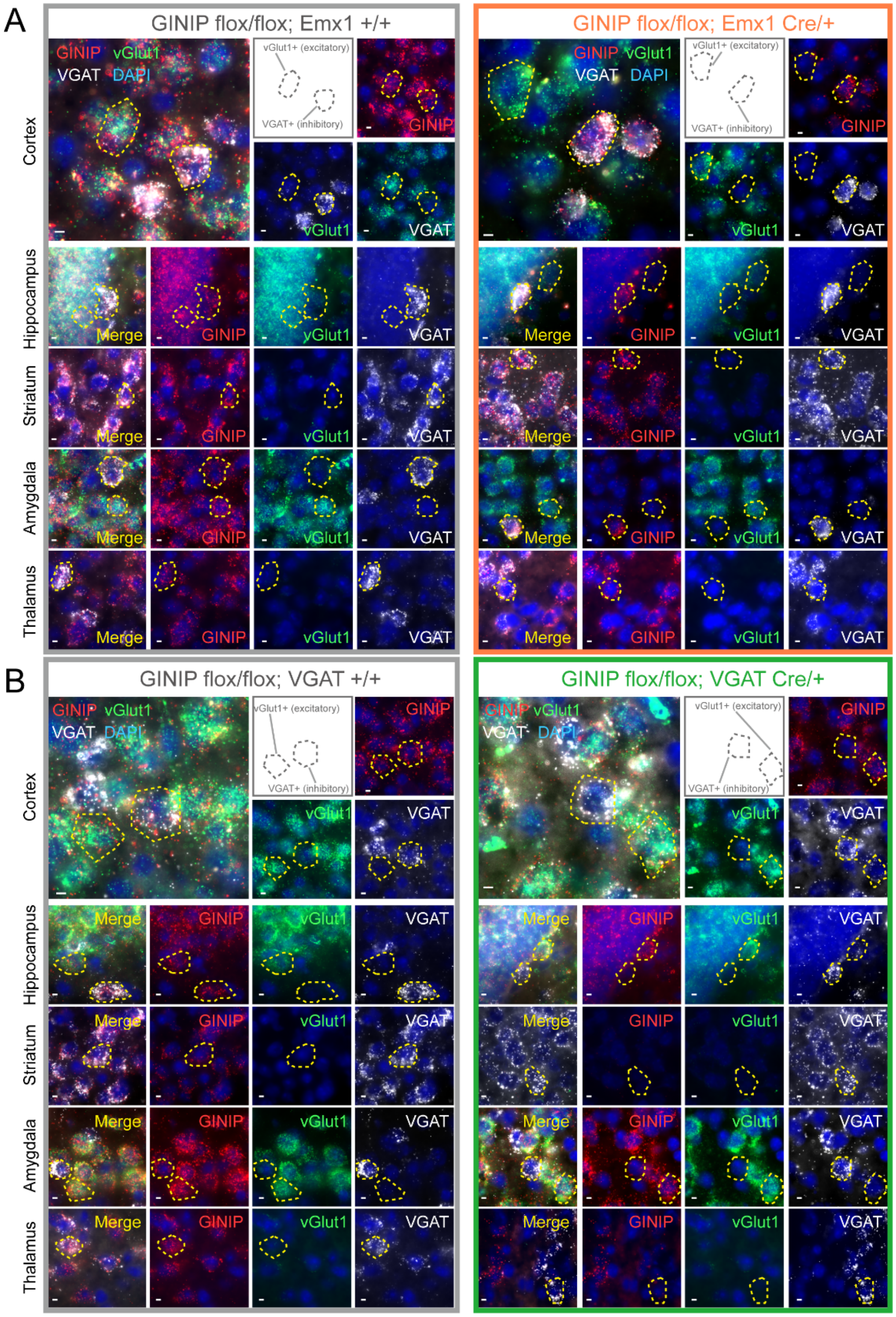


**Figure S6. GINIP expression is specifically ablated in excitatory (vGlut1$^{+}$) or inhibitory (VGAT$^{+}$) neurons of GINIP flox/flox mice upon expression of Cre in specific neuron populations.**

**(A)** GINIP expression is specifically ablated in excitatory (vGlut1$^{+}$) neurons, but not in inhibitory (VGAT$^{+}$) neurons, of GINIP flox/flox mice bearing an allele for Cre expression in Emx1$^{+}$ excitatory neurons. GINIP flox/flox mice were compared to littermates bearing an Emx1 Cre driver allele to achieve neuron-specific ablation of GINIP. GINIP, vGlut1 and VGAT mRNAs were simultaneously detected in the indicated regions of mouse brain slices by fluorescence in situ hybridization. Yellow dotted lines encircle the same cell across different panels of the same sample to facilitate the identification of GINIP$^+$ neurons that are either vGlut1$^+$ or VGAT$^+$.

**(B)** GINIP expression is specifically ablated in inhibitory (VGAT$^+$) neurons, but not in excitatory (vGlut1$^+$) neurons, of GINIP flox/flox mice bearing an allele for Cre expression in VGAT$^+$ inhibitory neurons. GINIP flox/flox mice were compared to littermates bearing an VGAT Cre driver allele to achieve neuron-specific ablation of GINIP. GINIP, vGlut1 and VGAT mRNAs were simultaneously detected in the indicated regions of mouse brain slices by fluorescence in situ hybridization. Yellow dotted lines encircle the same cell across different panels of the same sample to facilitate the identification of GINIP$^+$ neurons that are either vGlut1$^+$ or VGAT$^+$.

All scale bars are 10 μm. n = 3 for all results in this figure.
